# Expression and Purification of BsaXI Restriction Endonuclease and Engineering New Specificity from BsaXI Specificity (S) Subunit

**DOI:** 10.1101/2022.03.14.484276

**Authors:** Sonal Gidwani, Daniel Heiter, Shuang-yong Xu

## Abstract

BsaXI is a Type IIB restriction endonuclease (REase) that cleaves both sides of its recognition sequence 5’ ↓N9 AC N5 CTCC N10↓ 3’ (complement strand 5’ ↓N7 GGAG N5 GT N12↓ 3’), creating 3-base 3’ overhangs. Here we report the cloning and expression of *bsaXIS* and *bsaXIRM* genes in *E. coli*. BsaXI activity was successfully reconstituted by mixing the BsaXI RM fusion subunit with the BsaXI S subunit and the enzyme complex further purified by chromatography over 6 columns. As expected, the S subunit consisted of two subdomains encoding TRD1-CR1 (TRD, target recognition domain, CR, conserved region) for 5’ AC 3’, and TRD2-CR2 presumably specifying 5’ CTCC 3’. TRD1-CR1 (TRD2-CR2 deletion) or duplication of TRD1 (TRD1-CR1-TRD1-CR2) both generated a new specificity 5’ AC N5 GT 3’ when the S variants were complexed with the RM subunits. Circular permutation of TRD1 and TRD2, i.e. relocation of TRD2-CR2 to the N-terminus and TRD1-CR1 to the C-terminus generated the same specificity with the RM subunits, although some wobble cleavage was detected. The TRD2 domain in the BsaXI S subunit can be substituted by a close homolog (∼59% sequence identity) and generated the same specificity. However, TRD2-CR2 domain alone failed to express in *E. coli*, but CR1-TRD2-CR2 protein could be expressed and purified which showed partial nicking activity with the RM subunits. This work demonstrated that like Type I restriction systems, the S subunit of a Type IIB system could also be manipulated to create new specificities. Genome mining of BsaXI TRD2 homologs in GenBank found more than 36 orphan TRD2 homologs, implying that quite a few orphan TRD2s are present in microbial genomes that may be potentially paired with other TRDs to create new restriction specificities.

## Introduction

BsaXI is a Type IIB BcgI-like restriction endonuclease (REase) that cleaves outside of its recognition sequence 5’ ↓N9 AC N5 CTCC N10↓ 3’ (complement strand 5’ ↓N7 GGAG N5 GT N12↓ 3’), creating 3-base 3’ overhangs [1; 2]. It was originally found in the *Bacillus stearothermophilus* 25B strain (NEB catalog, www.neb.com; REBASE, new strain name *Geobacillus* sp. 25B) [3]. BcgI consisted of three subunits in the form of [RM]2 S in which the RM subunit is a fusion of endonuclease and N6mA methyltransferase (MTase) [4; 5]. It is also similar to a certain extent to Type IIG/IIB REases that form a single R-M-S fusion protein whose activity is sometimes stimulated by the presence of S-adenosylmethionine (SAM) [6]. However, some BcgI-like REase activities are independent of ATP and SAM. The domain organization of the S subunit of BcgI-like REases [4] is similar to the HsdS (S) subunit of Type I restriction systems (HsdR2/HsdM2/HsdS or (R2/M2/S)) in the linear order of TRD1-CR1-TRD2-CR2 (TRD, target recognition domain, CR, conserved region) [7; 8; 9]. In Type I HsdM2/HsdS complex structure, the CR1 and CR2 form two long α-helixes termed coiled-coil which interacts with the HsdM subunits to form a four-helix bundle and recognize two bipartite DNA sequences [10]. A structure model of Type I S subunit in complex with the R-M subunits has also been published previously, revealing two TRD domains of the HsdS subunit interacting with [HsdR/HsdM]2 subunits [11]. It has been shown that by site-directed mutagenesis of the S domain in the single chain MmeI-like endonucleases (R-M-S fusion), new specificities could be easily engineered by a few amino acid (aa) substitutions [12]. Since MmeI methylase domain shared the same specificity domain, the MTase target is also altered simultaneously with the restriction specificity. Specificity domain TRDs swapping among three Type IIG/B enzymes with related recognition sequences (AloI, PpiI, TstI) also created new restriction specificities [6]. Engineering new specificity from Type IIP REases proved to be more challenging [13; 14; 15; 16], unless creating relaxed activity (increased star activity) or shortened recognition sequences [17; 18]. However, by site-directed mutagenesis, one could isolate enzyme variants with significantly reduced star activity (“high-fidelity” HF mutants) of KpnI, BamHI, and EcoRI (SYX et al. unpublished results) [19] (US patents number 8,673,610 on BamHI-HF, US patent number 9,249,396 on EcoRI-HF). In some sequenced bacterial genomes, some HsdR and HsdM can potentially partner with a few specificity subunits to create multi-specificity Type I restriction systems as the HsdS genes are in the same operon as the R-M genes (REBASE).

Most of successful specificity (HsdS) engineering studies have been done with Type I restriction systems [20; 21]. In Type I REase EcoR124I, the N-terminal domain of the HsdS subunit (C-terminus deletion) can still interact with the HsdR2/HsdM2 subunits and generated a new specificity with symmetric recognition (WT EcoR124I (GAA N6 RTCG); EcoR124I HsdS_NT_ N-terminal domain (GAA N7 TTC)) [22; 23]. The length of a repeated sequences (two vs three copies) in the long α-helical region (CR1) of the S subunit in EcoR124I and Eco124/3 (EcoR124II) was shown to dictate the length of the non-specific DNA sequences between the two bipartite sites (GAA N6 RTCG vs. GAA N7 RTCG) [24]. Since the domain architecture of the S subunit of Type IIB restriction systems is analogous to that of Type I system, we hypothesize that truncation of TRDs in the S subunit may also create new specificity. Our goal is to define the functional domain of BsaXI TRDs and the boundary of TRD and CR, and the minimal length of CR sequences. In this work, we present the results of cloning and expression of BsaXI RM and S genes in *E. coli*. BsaXI enzyme complex can be purified by mixing cell extracts containing RM and S subunits or the RM and S subunits can be purified independently and two subunits can be mixed together to reconstitute BsaXI restriction activity. TRD1-CR1 was expressed and purified which was reconstituted with the purified RM subunits to create a new specificity (5’ AC N5 GT 3’). Part of the long α-helix in CR1 (42 aa) could be deleted: deletion of 8-aa, 15-aa, 21-aa, and 32-aa residues in CR1 created partial activity when the deletion variants were complexed with RM subunits. TRD1 duplication in TRD1-CR1-TRD1-CR2 also showed partial activity. We also demonstrated that CR1 can be replaced by CR2 in TRD1-CR2 and generated a partial activity. Circular permutation of BsaXI TRDs in TRD2-CR2-TRD1-CR1 (i.e. relocation of TRD2 to the N-terminus and TRD1 to the C-terminus) produced the same specificity as the WT enzyme, although cleavage at the 3’ side of CTCC and GT is somewhat imprecise and wobbly (5’ ↓N9 AC N5 CTCC N10-13↓ 3’ (complement strand 5’ ↓N7 GGAG N5 GT N12-13↓ 3’). Soluble TRD2-CR2 protein could not be expressed in *E. coli*. But CR1-TRD2-CR2 variant protein could be expressed and purified, which showed low DNA nicking activity when it is reconstituted with the RM subunits. In addition, we showed that the BsaXI TRD2 can be substituted by a close homolog (59% aa sequence identity) and the chimeric S subunit created the same specificity as the WT enzyme. However, substitution with a distant TRD2 homolog failed to produce active S variant. Protein homolog search using BlastP in GenBank found very few standalone BsaXI TRD1 homologs in microbial genomes, but there are at least 36 orphan BsaXI TRD2 homologs, implying TRD2-CR2 in bacteria are probably available to pair up with other functional TRDs to create new restriction systems. This work provided more understanding of the S subunit of Type IIB restriction system and demonstrated a direct strategy to engineer new specificity from Type IIB systems.

## Materials and Method

Restriction enzymes, 2x Phusion DNA polymerase PCR master mix, T4 DNA ligase, pBR322, pUC19, pTYB1, chitin beads and Gibson DNA assembly/cloning kit were supplied by New England Biolabs (NEB). Bacterial genomic DNA (gDNA) was purified from *Bacillus stearothermophilus* 25B strain (NEB strain collection, new strain name *Geobacillus* sp. 25B) by a phenol-chloroform extraction method as described before [25]. Big-Dye Sanger sequencing kit was purchased from Thermo-Fisher/ABI and plasmid DNA was sequenced with the protocol recommended by the manufacturer. The amplified PCR fragments of *bsaXIRM* and *bsaXIS* genes were cloned into intein-CBD expression vector pTYB1 (NdeI-XhoI cut, NEB) [26] and the inserts were sequenced to verify the correct coding sequences. The plasmids were transferred into T7 expression host C2566 (NEB) by transformation. DNAStar (Lasergene 14) and Geneious softwares were used for DNA sequence editing.

### BsaXI enzyme complex purification

C2566 [pTYB1-*bsaXIRM*] and C2566 [pTYB1-*bsaXIS*] cells were cultured in 1-liter (L) flasks of LB + ampicillin and grown at 30°C until OD600 reached 1.3. After sitting at room temperature for 30 min, cultures were induced with 0.4 mM IPTG and grown overnight at 18°C. Cells were harvested by centrifugation and stored at -80°C. Frozen cell pellets from 3 L of BsaXI RM culture and 6 L of BsaXI S culture were all thawed (preliminary experiment indicated that the expression level of RM subunit is higher than the S subunit), resuspended in chitin column buffer (500 mM NaCl, 20 mM Tri-HCl, pH 8.5, 1 mM EDTA, and 0.1% Triton X-100 or Tween 20), lysed through 2 passes at 30 kpsi in a Dyhydromatics HL60 cell disruptor, and centrifuged to remove cell debris. Crude supernatants were loaded through two separate 40 ml chitin resin (NEB S6651) columns by gravity, washed with 300 ml of chitin column buffer, and then cleaved over 3 days with 50 mM DTT chitin cleavage/elution buffer. All chitin eluates were pooled together in 450 ml, diluted to 300 mM NaCl with addition of 300 mL zero salt column buffer and flowed through 20 ml DEAE resin to remove nucleic acids. DEAE flow-through and wash were again diluted to 150 mM NaCl with zero salt column buffer, applied to a Heparin TSK column (11 ml) and eluted with increasing NaCl gradient. Pooled active fractions were dialyzed overnight against 100 mM NaCl, 20 mM Tris-HCl, pH 8, 1 mM DTT, 0.1 mM EDTA, and 5% glycerol, passed through a Source 15S column (21 ml), then applied to 22 ml Source 15Q and eluted with increasing NaCl gradient (0.1 to 1 M). Pooled active fractions were dialyzed overnight against 50 % glycerol (NEB Diluent B) and applied to a Superdex75 SEC column (1787 ml). Pooled active fractions were concentrated by a second run over Heparin TSK. Final yield was ∼10.0 mg pure BsaXI REase ([RM]2 + S). One BsaXI unit (U) is defined as the amount of protein required for complete digestion of 1 μg phage λ DNA in CutSmart buffer at 37°C for 1 h.

### Separate purification of BsaXI RM and S subunits

2 L of IPTG-induced cells containing RM-intein-CBD or 6 L of IPTG-induced cells of S-intein-CBD were resuspended in 60 ml and 180 ml of chitin column buffer and the suspensions sonicated to lyse the cells. After removal of cell debris by centrifugation at 15k rpm at 4°C for 30 min, clarified cell lysates were loaded onto 20 ml chitin columns by gravity flow (1 column for RM-intein-CBD, 2 columns for S-intein-CBD). CBD-tagged enzymes in flow-through were reloaded three times and the columns were washed with 10 column volumes of chitin buffer (∼200 ml). 20 ml of cleavage buffer (chitin column buffer + 50 mM DTT) was added to the top of chitin column. Approximately 2 ml of cleavage buffer passed through the column and the flow-through was discarded. The DTT-catalyzed intein cleavage reaction continued at 4°C for 2-3 days. The eluted proteins from chitin columns were diluted in a low salt buffer to reach 0.1 M NaCl concentration, which was subsequently loaded onto 5 ml Hi-Trap Heparin column (GE Healthcare). After extensive washing with low salt, the RM or S protein was eluted with a salt gradient (0.1 – 1 M NaCl). Eluted peak fractions were analyzed by SDS-PAGE, concentrated in protein concentrators and protein was resuspended in enzyme storage buffer to be stored at -20°C.

### BsaXI Specificity (S) subunit domain organization and amino acid (aa) sequence of S subunit variants

Synthetic genes (gene blocks) encoding S variants were purchased from IDT, which carry 24-27 bp overlapping vector sequences at the 5’ and 3’ ends. Synthetic genes were assembled into pTYB1 using Gibson assembly kit and the assembled DNA was transferred into T7 expression strain C2566 (T7 Express, NEB) by transformation. To construct 6xHis-tagged S expression clone, A PCR fragment containing *bsaXIS* gene was cloned into pET21b (NdeI and XhoI cut, Novagen) to achieve C-terminal 6xHis tag of the targe protein. pET21-*bsaXIS* plasmid was transferred into T7 Express (C2566) for expression. BsaXI S (6xHis) tagged protein was purified from a nickel agarose beads column (10 ml, NEB) by gravity flow using a protocol supplied by NEB.

WT: TRD1-CR1-TRD2-CR2 (54.98 kDa)

TRD1 and TRD2: aa residues shown in blue and green, respectively. CR1 and CR2, underlined aa residues. The boundary of CR1 (42-aa) and CR2 (42-aa) are predicted by structured-guided Phyre2 aa sequence alignment [27]. PIPP residues shown in red indicates the boundary between TRD2 and CR2 (see **suppl. Fig. S1** for sequence alignment). The boundary between TRD1 and CR1 is less clear-cut as CR1 may include 1-2 additional aa residues depending on the secondary structure prediction software; similarly the boundary between CR1 and TRD2 has a 5-aa sequence (HFNIN) that is not part of either domain (see suppl. Fig. S1).

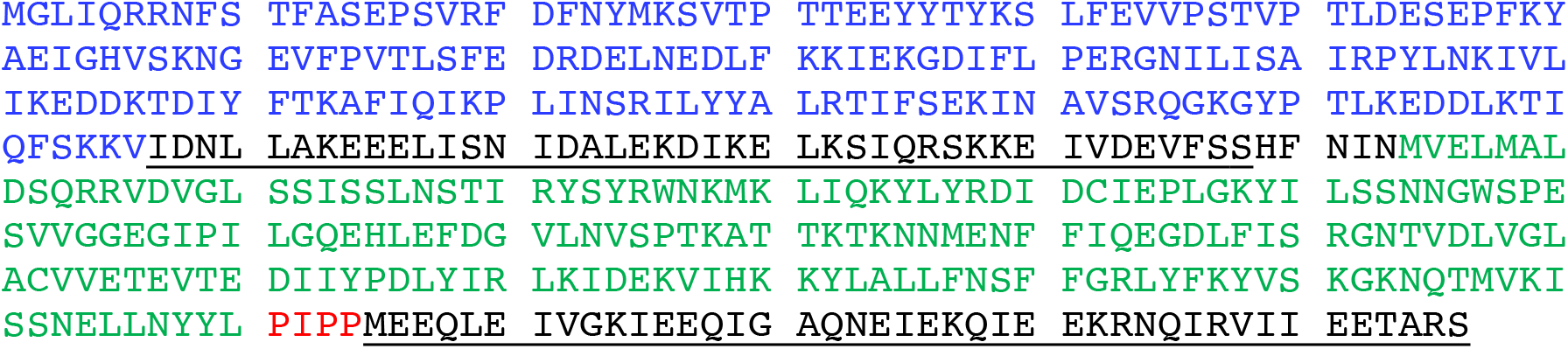

TRD1-CR1 (TRD2-CR2 deletion, 27.06 kDa)

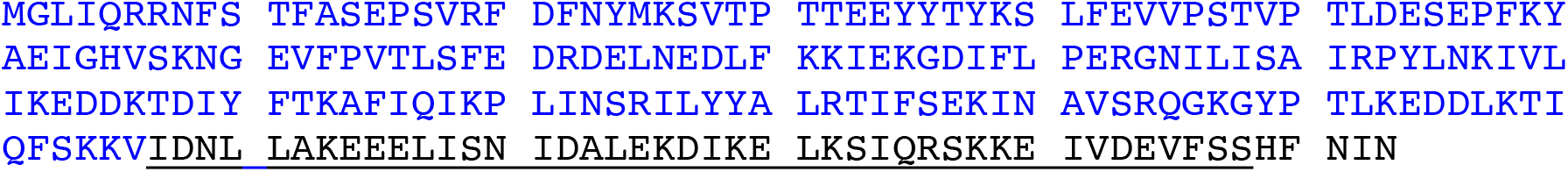

TRD1-CR2 (TRD2 deletion, replacing CR1 with CR2, 27.53 kDa)

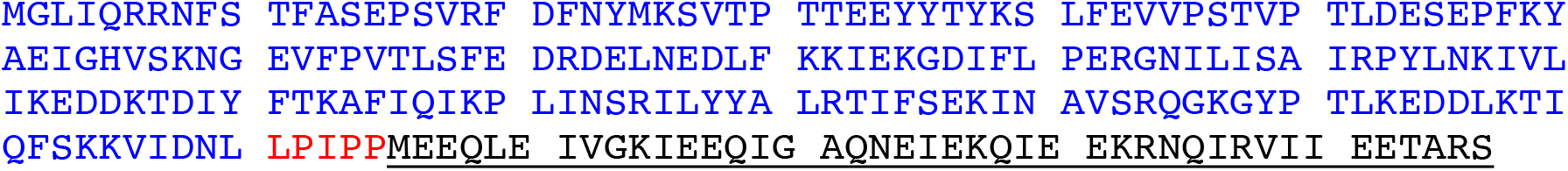

TRD1-CR1-8aaΔ (8-aa deletion in the predicted long α-helix at the C-terminus, 25.56 kDa)

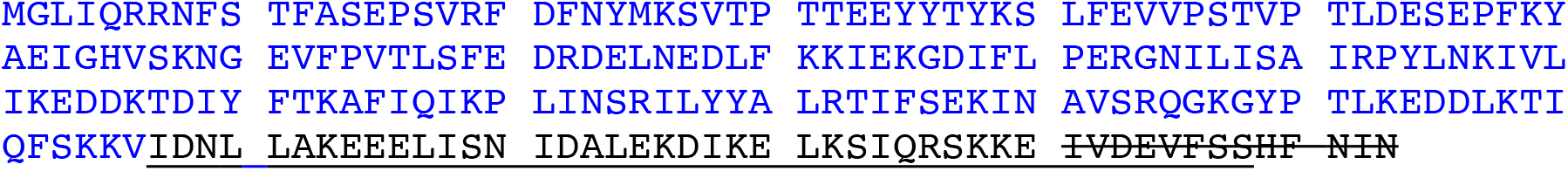

TRD1-CR1-15aaΔ (15-aa deletion in the predicted long α-helix at the C-terminus, 24.69 kDa)

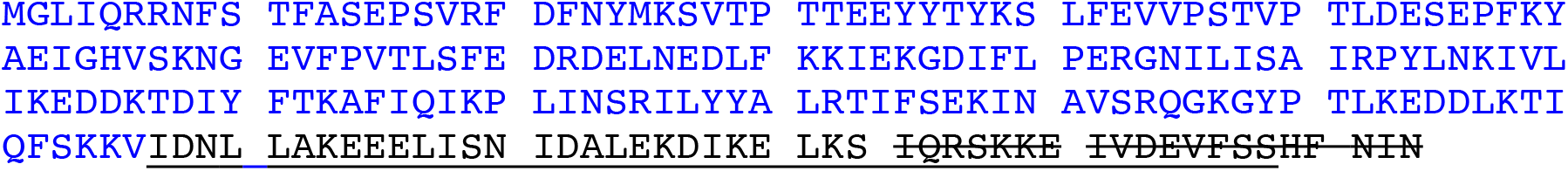

TRD1-CR1-21aaΔ (21-aa deletion in the predicted long α-helix at the C-terminus, 23.99 kDa)

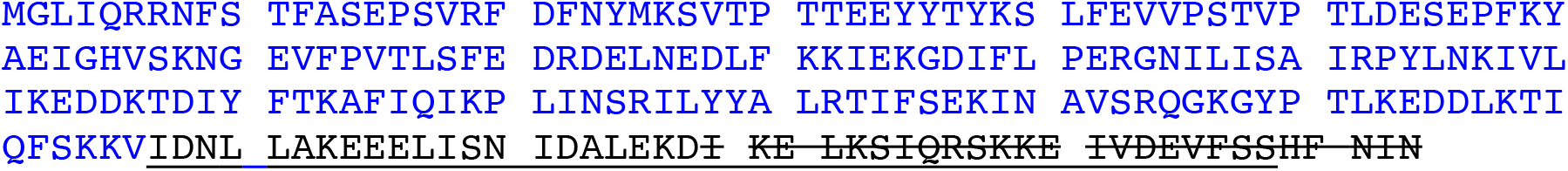

TRD1-CR1-32aaΔ (32-aa deletion in the predicted long α-helix at the C-terminus, 22.78 kDa)

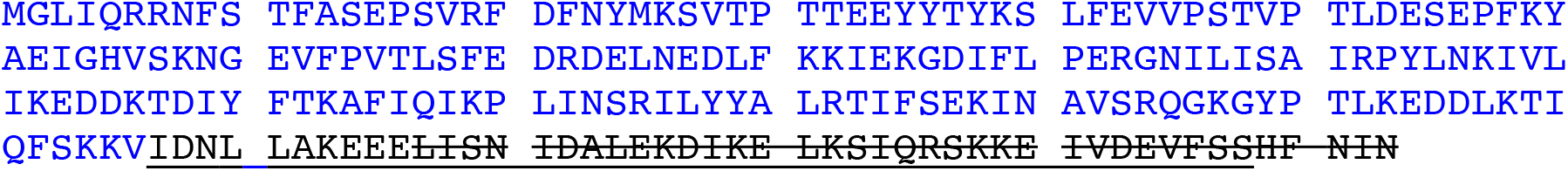

TRD1-CR1-57aaΔ (15-aa deletion in TRD1 plus 42-aa deletion of the entire predicted long α-helix at the C-terminus, protein could not be expressed in *E. coli*)

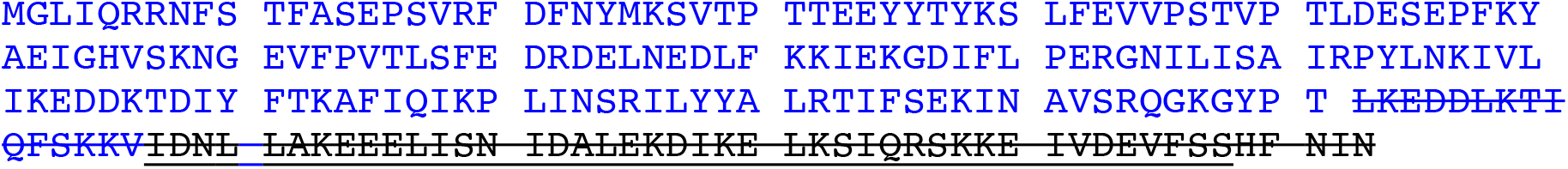

TRD1-CR1-TRD1-CR2 (TRD1 duplication, a.k.a., 2xTRD1 with CR1 and CR2, 54.11 kDa)

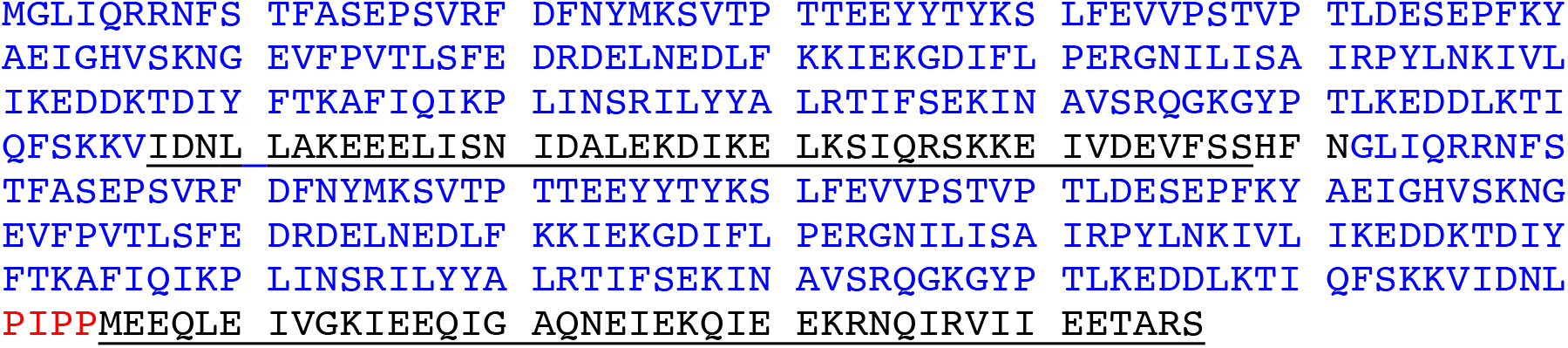

TRD2-CR2 (ΔTRD1-CR1, protein could not be expressed in *E. coli*)

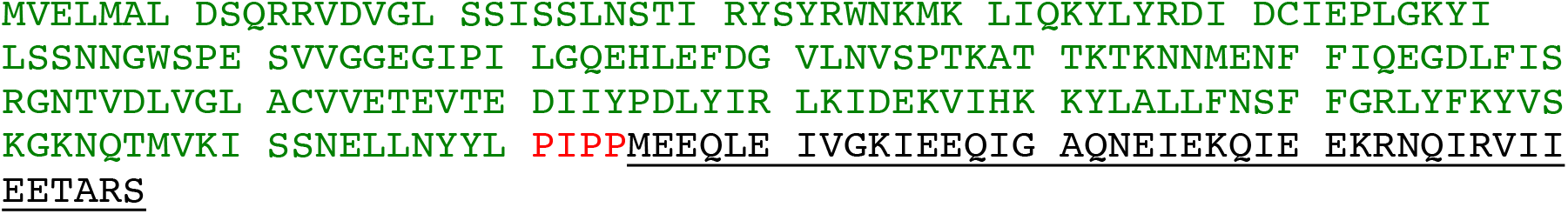

CR1-TRD2-CR2 (ΔTRD1, 33.16 kDa)

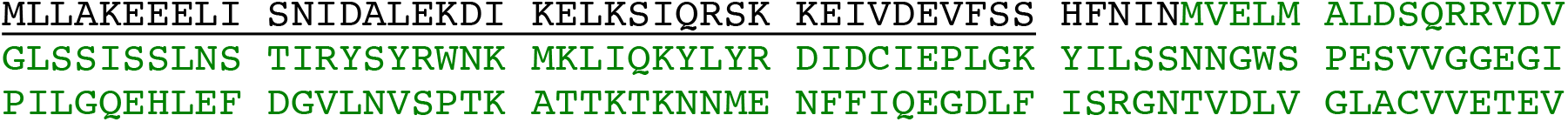
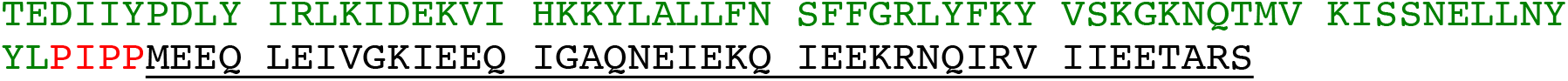

### Circular permutation of TRDs

rearranged TRD domains: TRD2-CR2-TRD1-CR1 (54.85 kDa)

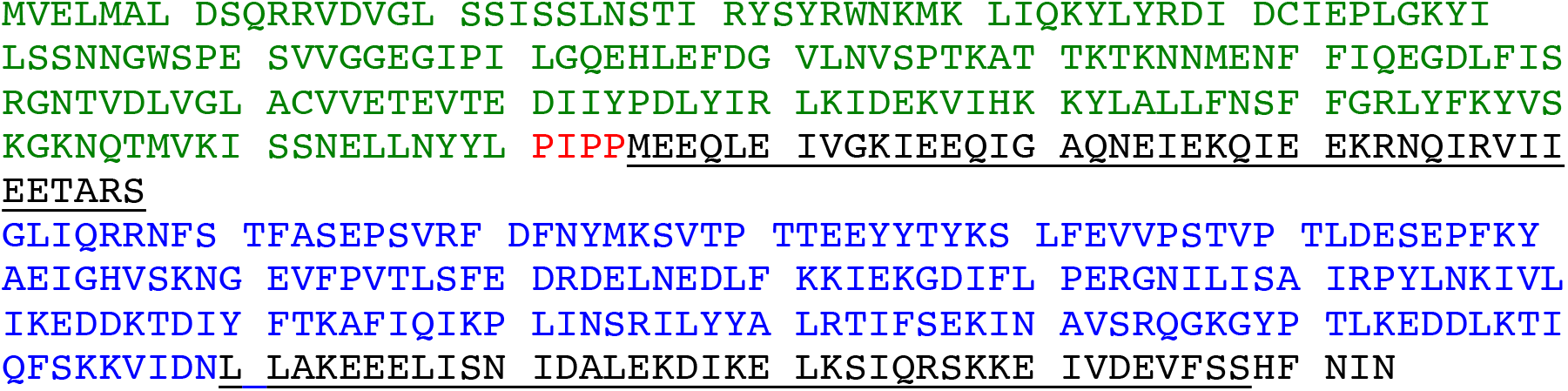

### Replacing BsaXI TRD2 with a TRD homolog from another bacterium

CspC0110I TRD homolog (partial) was found in GenBank by BlastP search with 59% aa sequence identity to BsaXI TRD2 (see below for GenBank accession number). It was annotated as a Type I HsdS partial sequence in *Cyanothece* sp. CCY0110.

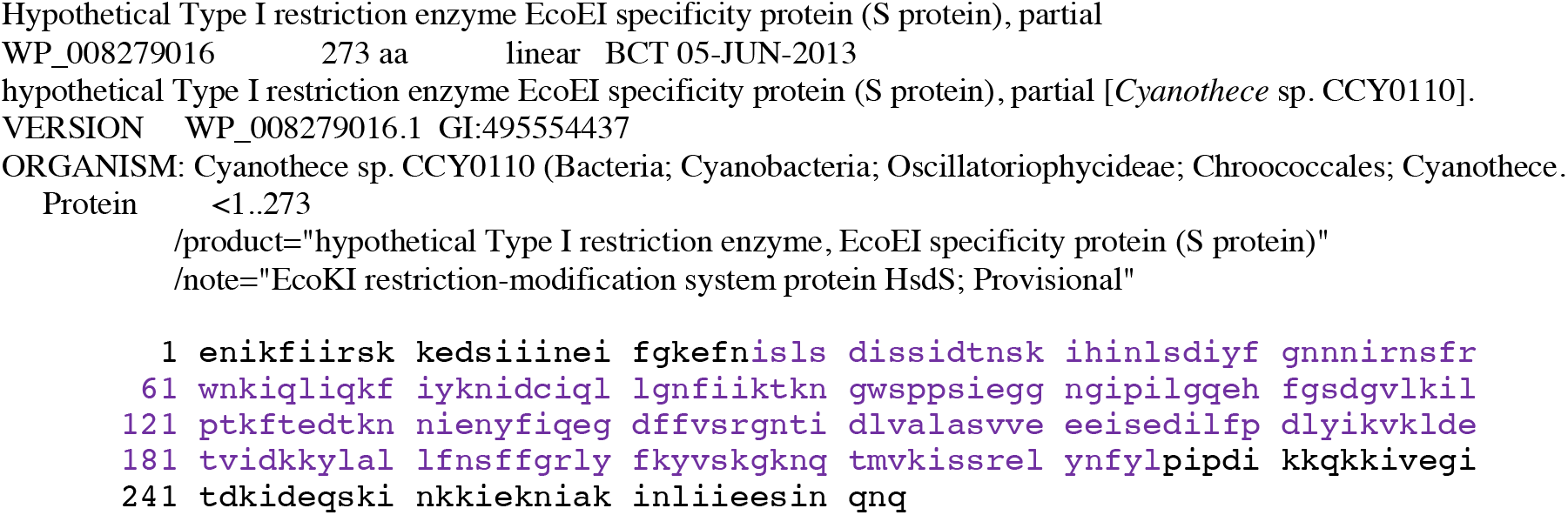

TRD1-CR1-[CspC0110I TRD]-CR2 (55.04 kDa)

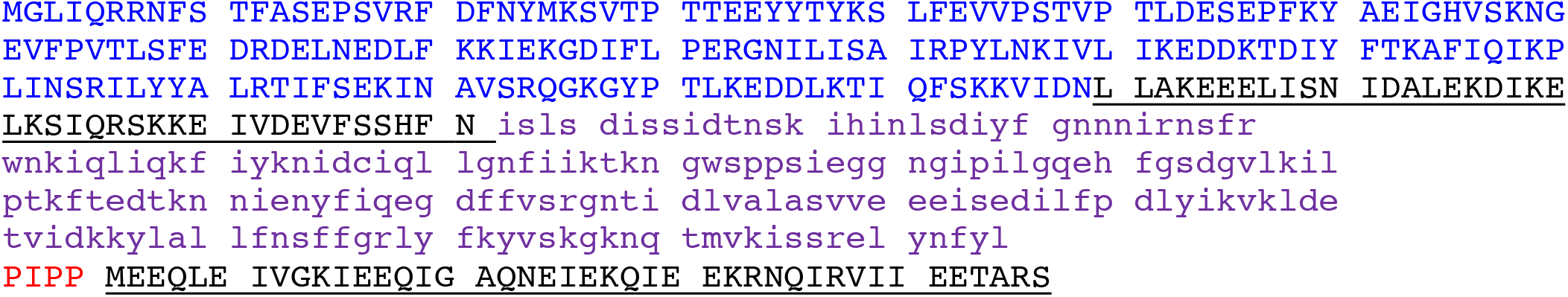

Chimeric S subunit TRD1-CR1-[Cst TRD]-CR2

BsaXI TRD2 and Cst TRD2 share 28% aa sequence identity. The Cst S subunit is annotated as a Type I specificity protein in GenBank (accession number WP_005530828).

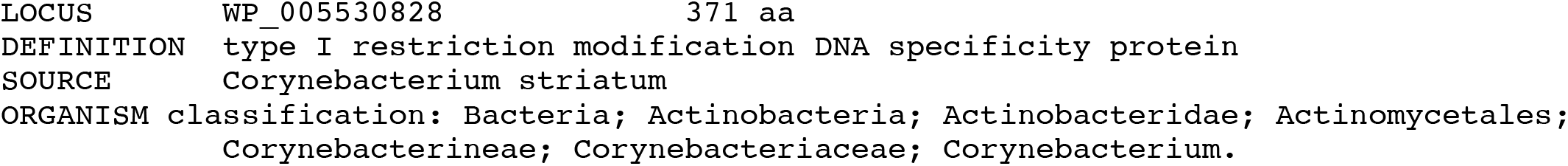

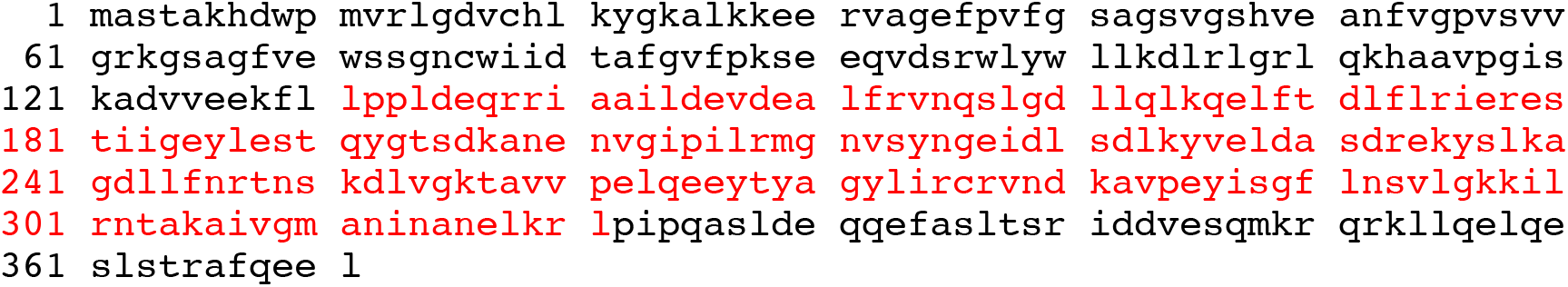

Chimeric S subunit: TRD1-CR1-[Cst TRD]-CR2 (no protein expression)

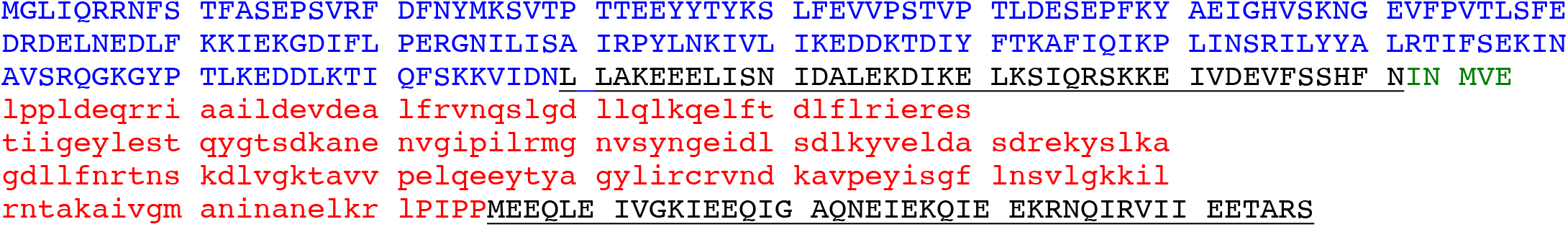

## Results

### Cloning of BsaXI restriction-modification genes in *E. coli*

The BsaXI restriction-modification (R-M) system was originally identified from random shotgun sequencing of *B. stearothermophilus* 25B (*Geobacillus* sp. 25B) genomic DNA (data not shown). Part of the sequences were verified by primer walking using Sanger sequencing. The R-M system contains two genes, the *bsaXIRM* gene encoding the restriction-modification subunit (RM fusion, 911 aa, predicted molecular mass 107.04 kDa) and the *bsaXIS* gene coding for the specificity subunit (S, 476 aa, molecular mass 54.97 kDa). The RM and S genes are probably regulated in a single transcription unit with RM gene preceding the S gene. The DNA sequence has been deposited in GenBank (accession number OM373208). The endonuclease domain in the N-terminal region of RM subunit contains two potential PD-D/ExK catalytic sites (D-X16-DxK or D-X17-ExK). It is predicted that the second motif may form the catalytic site since it is analogous to the catalytic residues of BcgI: PE-X12-ExK [4]. The exact catalytic residues remain to be confirmed experimentally. The methylase domain in the C-terminal region contains typical N6mA (6mA) methyltransferase (MTase) conserved sequence motifs. The methylase activity is predicted to modify the adenine (A) in the target site 5’ ↓N9 **A**C N5 C**<u>T</u>**CC N10↓ 3’, and the adenine in the bottom strand opposite to the <u>T</u> base. The MTase activity of the purified BsaXI RM-S enzyme complex is low and only provided partial modification in vitro (see below).

Sequence alignment with the S subunits of other Type I and Type IIB systems by Phyre2 indicated that the S subunit contains protein subdomains TRD1-CR1-TRD2-CR2 in which CR1 and CR2 are predicted to form long α—helix coiled-coil and interact with the RM subunits (see below for more detailed analysis of TRDs, **suppl. Fig. S1**). BsaXI RM and S gene products are nearly identical to two putative R-M systems listed in REBASE: 99% to 100% sequence identity to Gth3921I and Gka8005I (RM fusion, WP_052369193; S subunit, WP_052369191; *Geobacillus kaustophilus* NBRC 102445 DNA sequence contig, NZ_BBJV01000045), suggesting that the BsaXI R-M system (isoschizomers) may have been evolved in other *Geobacillus* strains via horizontal gene transfer. Homologs of BsaXI R-M systems are widely distributed in other microbial genomes: more than 60 RM fusion homologs with 44% to 100% aa sequence identity are found in GenBank in a BlastP search (data not shown).

### Expression and purification of BsaXI restriction endonuclease

Co-expression of BsaXI RM and S genes in pUC19 appeared to be unstable (data not shown). The reason for this might be insufficient methylation. Therefore, the two genes were expressed separately in fusion with intein and chitin binding domain (CBD) in pTYB1. The fusion to intein and CBD may also contribute to the reduced toxicity of RM expression. Cell extracts containing RM and S subunits could be mixed in test tube and the RM/S subunits in the complex was co-purified. **Figure 1A** shows the co-purified BsaXI after chromatography through six columns (Chitin, DEAE, Heparin Sepharose, Source 15 S, Source 15Q, and gel filtration). Based on the protein molecular mass and band intensity (Figure 1A and suppl. Fig. S2) it was estimated that enzyme stoichiometry of RM : S is 2 : 1 (band intensity (total pixels ratio) = 2 : 1.2 as measured by gel imaging software, Bio Rad), and the active enzyme complex can be shown as trimeric complex [RM]2 S similar to the first characterized Type IIB enzyme BcgI [4]. However, we cannot rule out the possibility of forming dimer of trimer ([RM]2 S)2 in the presence of cognate DNA. Two BsaXI subunits could be purified separately by chromatography through chitin and heparin columns (**Figure 1B**). To increase the expression level for BsaXI S subunit, we also constructed an expression clone in pET21b with 6xHis tag. BsaXI S (6xHis) was purified by chromatography through a nickel agarose column (**suppl. Fig. S2**). **Figure 2A** shows the restriction activity of co-purified BsaXI endonuclease. At high enzyme concentrations the activity was inhibited either due to protein aggregation or to a lack of available target sites as “overcrowded” enzyme non-specific binding. BsaXI activity can also be reconstituted by mixing the partially purified RM and S (6xHis) subunits to achieve partial digestion (**Figure 2B**).

**Figure 1.**
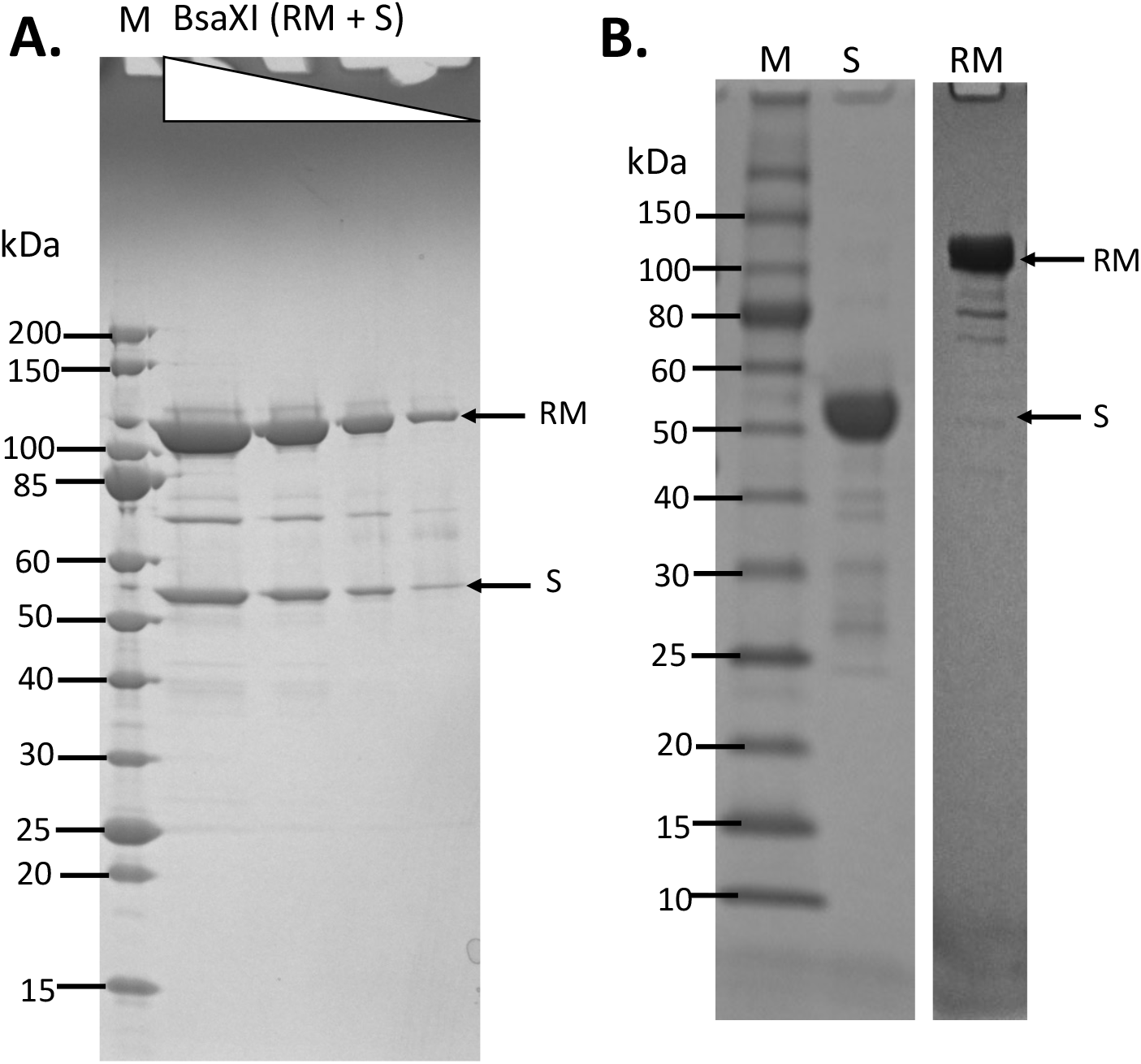
Purification of BsaXI RM and S subunits and BsaXI enzyme complex. **A**. BsaXI enzyme complex (RM + S) purified by 6-steps chromatography (see details in Materials and Method). BsaXI was analyzed on SDS-PAGE by 3-fold serial dilutions. The predicted molecular mass of the RM and S subunits are 107 and 55 kDa, respectively. M, protein molecular mass ladder (NEB). **B**. SDS-PAGE analysis of partially purified BsaXI S and RM subunits. BsaXI RM and S subunits were purified separately by chromatography through chitin and heparin columns.

**Figure 2.**
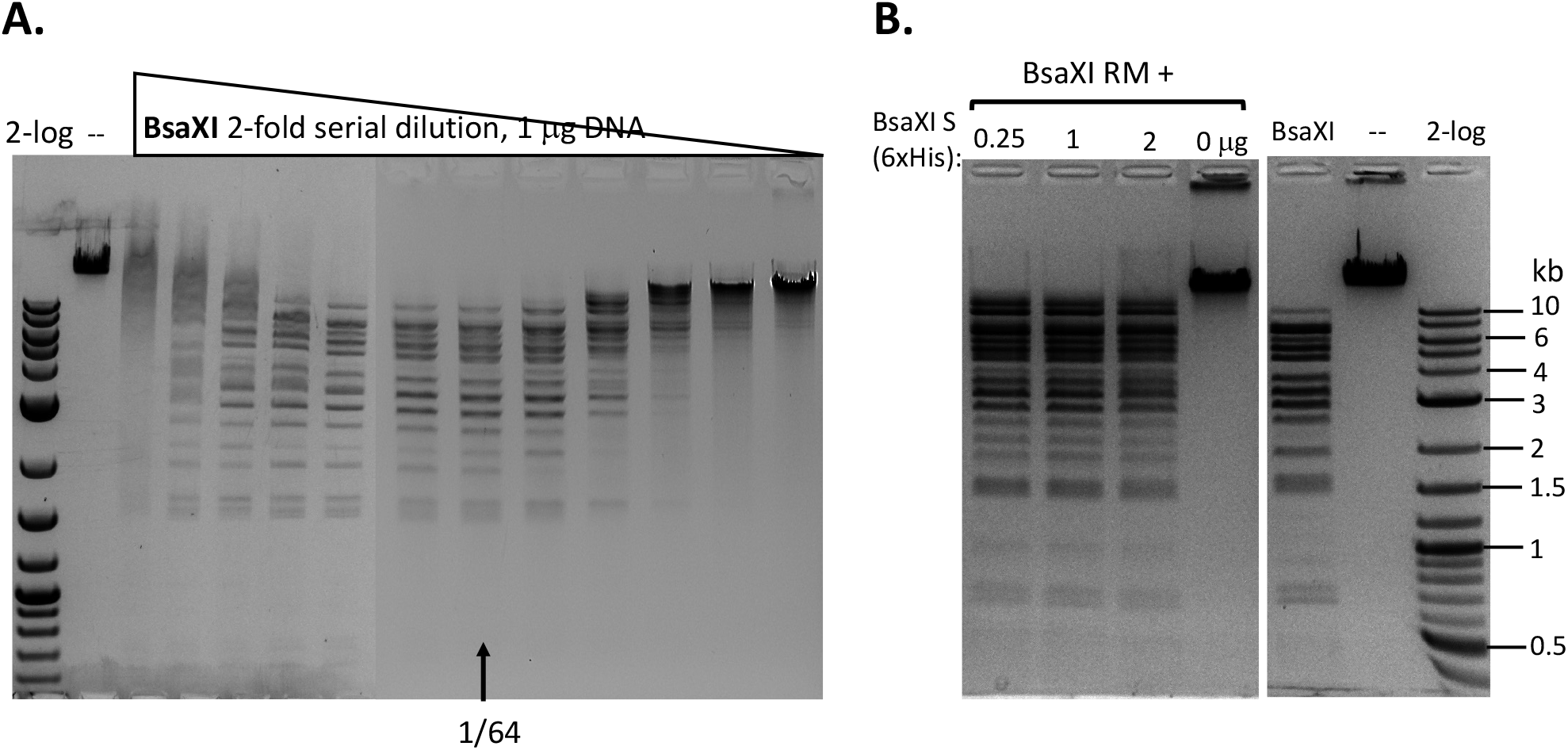
Recombinant BsaXI activity assays. **A**. BsaXI restriction activity by co-purified BsaXI enzyme (RM+S). The specific activity of BsaXI was estimated at 22,000 U/mg protein (see unit definition in Materials and Method). A complete digestion pattern was observed at 1/64 enzyme dilutions. At high enzyme concentration, the DNA was strongly bound and shifted upwards, suggesting enzyme aggregation with the DNA. **B**. BsaXI restriction activity reconstituted by mixing purified RM (chitin/DEAE/Heparin columns) and S (6xHis) (Ni agarose column) subunits in restriction digestion of λ DNA. Fixed amount of RM subunit (1 μg, at ∼93.5 nM, lanes 1-3) was mixed with varying amount of S (6xHis) (0.25, 1, and 2 μg, at 90.9, 363.6, and 727.2 nM). RM to S ratio approximately at 1/1, ¼, and 1/8 in lanes 1-3; lane 4, BsaXI RM subunit only; lane 5, BsaXI positive control (4 U); lane 6, uncut DNA; Lane 7, 2-log, DNA size ladder (0.1 to 10 kb, NEB).

Increasing BsaXI RM subunit concentration slightly improved the cleavage reaction, but still showing partial digestion (**suppl. Fig. S3**). The activity of in vitro reconstituted BsaXI activity by mixing the two subunits together in digestion of λ DNA is lower than the co-purified enzyme complex, probably due to slow subunit unit association, non-optimal subunit ratio or non-productive complex formation.

### Alteration and rearrangement of BsaXI S subunit

#### TRD1-CR1 (deletion of TRD2-CR2), TRD1-CR1-8aaΔ, TRD1-CR1-15aaΔ, TRD1-CR1-21 aaΔ, TRD1-CR1-57aaΔ

The C-terminal region of the S subunit (TRD2-CR2) was deleted to construct an S variant TRD1-CR1. Synthetic gene encoding TRD1-CR1 was cloned into pTYB1 and the insert was verified by Sanger sequencing. Following IPTG induction the protein was purified from a chitin column (**suppl. Fig. S2**) and its endonuclease activity was reconstituted with purified RM subunits. No BsaXI sites are present in pBR322 so any cleavage of the substrate by the S variant/RM should be a new cleavage specificity. pBR322 was digested into small fragments of less than 1 kb as detected in the agarose gel in **Figure 3A** (lanes 1-2). TRD1-CR deletion variants with 8-32 aa deleted also showed partial activity (**Figure 3B**, lanes 3-8). TRD1-CR1 with 21-aa deletion is also partially active (data not shown). **Figure 3C** shows the virtual restriction digestion of pBR322 by NEBcutter [28] on predicted 5’ AC N5 GT 3’ sites. The digested DNAs were subjected to run-off sequencing and the cut sites were found N9-12 bases outside of recognition sequence 5’ AC N5 GT 3’ (**suppl. Fig. S4A, B**). Cleavages at more than 12 nt distance were also detected in TRD1-CR1/RM digested DNA, but it is not clear whether the wobble cleavage is carried out by the TRD1-CR1/RM complex or by a contaminating exonuclease or cleavage directed from star sites. We also detected a cleavage outside of a star site at 5’ TC N5 GT N9↓ 3’ (**suppl. Fig. S4C**). It was concluded that TRD1-CR1 in complex with BsaXI RM subunit created a new restriction specificity (AC N5 GT) and cleavage took place outside of its recognition sequence. Part of CR1 in the long α-helix in TRD1-CR1 can be deleted up to 32-aa residues, but the deletion variants in CR1 region impaired the endonuclease activity. It was somewhat unexpected that large deletion in the CR1 region did not alter the spacer N5 in AC N5 GT. We also attempted to delete the entire CR1 region, but an expression clone with 42-aa deletion could not be constructed. Deletion of the entire CR1 plus 15-aa residues in the TRD1 domain failed to express the mutant protein, suggesting the 57-aa deletion negatively affected protein folding and expression.

**Figure 3.**
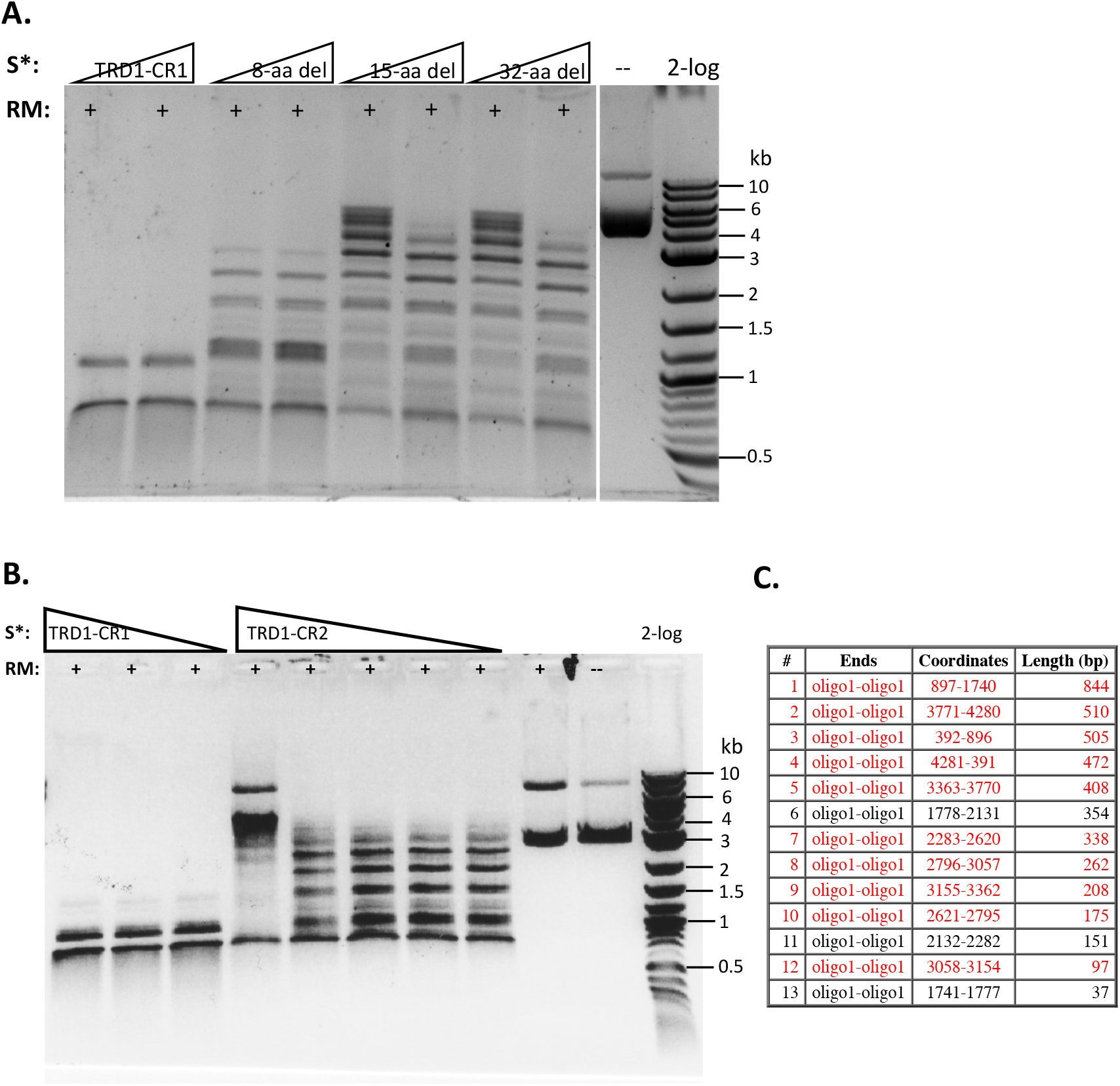
Reconstitution of restriction activity by mixing WT RM subunits with S variants. **A**. DNA cleavage patterns of pBR322 generated by reconstitution of BsaXI RM subunit with S variants TRD1-CR1 or TRD1-CR1 deletion variants (8-aa, 15-aa, 32-aa deletions in CR1). Fixed amount of RM subunit (1 μg, 94 nM) was mixed with two concentrations of the S variants (185 and 370 nM of 8-aa, 15-aa, 32-aa deletions in CR1, respectively). **B**. DNA digestion patterns of pBR322 by reconstitution of BsaXI RM (1 μg, 94 nM) subunit with S variants TRD1-CR1 (370, 185, and 93 nM, lanes 1-3), TRD1-CR2 (CR2 replacing CR1) (1450, 725, 363, 181, and 91 nM, lanes 4-8). High concentration of TRD1-CR2 inhibited activity (lane 4). Digestions were carried out in 1x CutSmart buffer at 37°C for 1 h. No BsaXI sites exist in pBR322, so observed cleavage suggested a new restriction specificity. Lane 9, RM subunit only. Lane 10, uncut DNA. **C**. Computer generated virtual cleavage of pBR322 in the sites 5’ AC N5 GT 3’ by NEBcutter.

#### TRD1-CR2 (deletion of TRD2 and replacing CR1 by CR2)

We also constructed and purified TRD1-CR2 (CR1 was replaced by CR2). This S variant generated partial restriction activity on pBR322 in complex with the RM subunits (**Figure 3B**, lanes 4-8). It was concluded that the long α-helix forming CR2 can replace CR1 and TRD1-CR2 can still form active complex with the RM subunits to create limited partial activity. It would be interesting to swap in other related CRs in Type I and IIB S subunits and test their interactions with BsaXI RM subunits. It was concluded that CR1 sequence is not absolutely required for TRD1 to interact with the RM subunits. Partial deletion of the CR1 segment did not completely eliminate TRD1 interactions with the RM subunits (see below).

#### TRD1 duplication: TRD1-CR1-TRD1-CR2 (deletion of TRD2)

We also attempted expression and purification of 2xTRD1 (TRD1-CR1-TRD1-CR2) in which TRD2 was replaced by TRD1. The S variant was purified from a chitin column and restriction activity was reconstituted with purified RM subunits. The duplicated TRD1 S variant generated partial restriction activity (data not shown). The digested DNA was subjected to DNA run-off sequencing and the recognition sequence was confirmed to be 5’ AC N5 GT 3’, but the cut sites are variable. When a DNA template strand is cut (nicked), the sequencing Taq DNA polymerase adds an extra peak “A” (adenine) to the sequencing read after the broken backbone (i.e. template independent terminal nucleotide transferase activity), creating doublets such as A/T, A/C, or A/G. If the original base call is “A”, then the extra overlapping “A” will create a high “A” base call. The sequencing peaks after the broken template will significantly drop off, thus termed DNA run-off sequencing. In a partially digested template, some DNA molecules are cut/nicked, while others remain intact and the base calls sometimes continue after the sudden peak drop off. Three examples of cut sites are shown in **Figure 4**. The cleavage took place either upstream or downstream as asymmetric cleavage AC N5 GT N9-12, which differs from coordinated WT cleavages on both 5’ and 3’ of its recognition sequence 5’ AC N5 CTCC 3’. We concluded from this experiment that 2xTRD1 (TRD1-CR1-TRD1-CR2) can interact with the RM subunits and activate cleavage, although cleavage near the symmetric sequence 5’ AC N5 GT 3’ appeared to be asymmetric.

**Figure 4.**
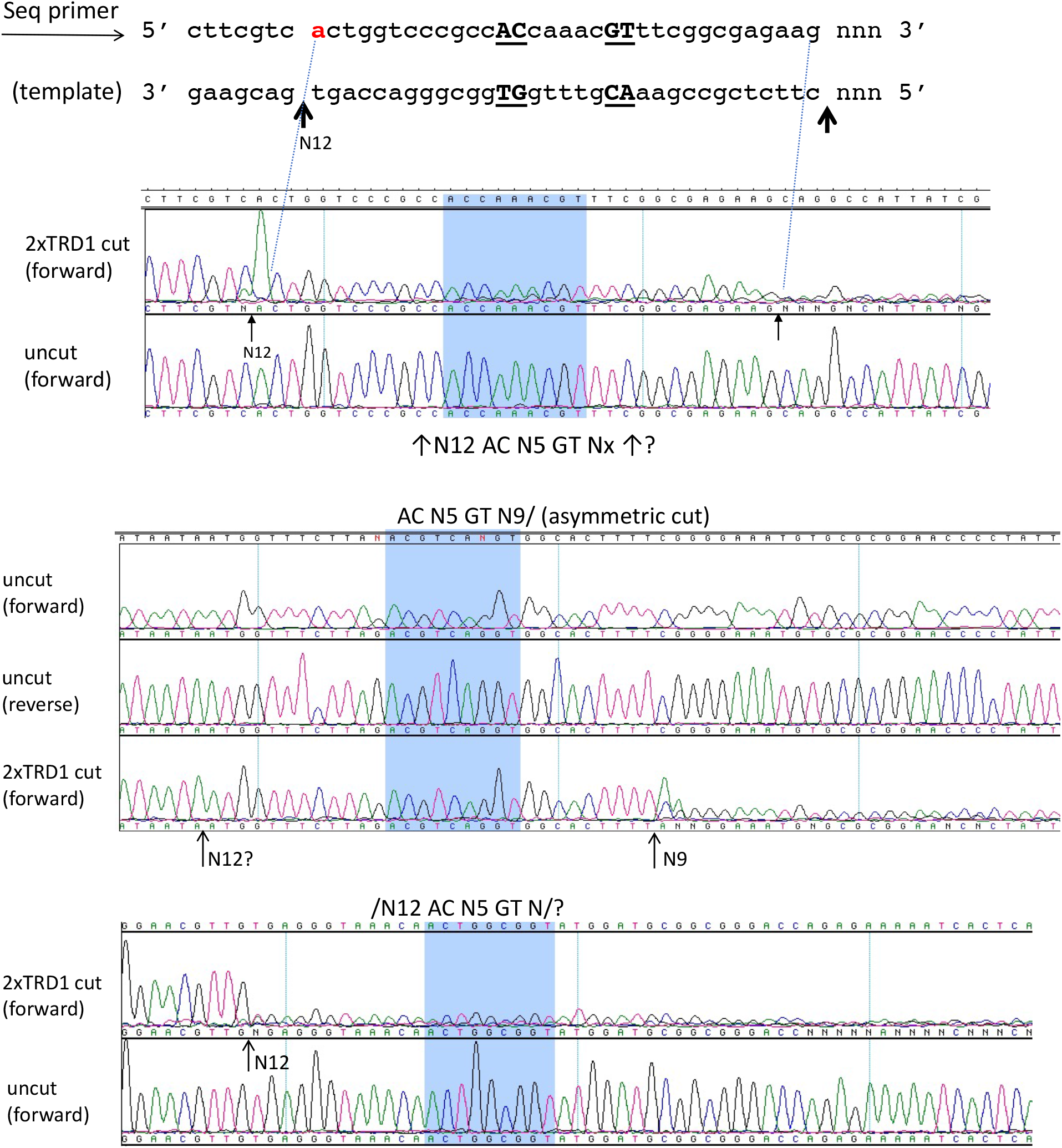
DNA run-off sequencing of 2xTRD1 (TRD1-CR1-TRD1-CR2)/RM digested pBR322 DNA to show the cut sites outside of 5’ AC N5 GT 3’. When the template strand is cut, the sequencing Taq DNA polymerase adds an extra peak “A” (adenine) to the sequencing read after the broken backbone, creating doublets such as A/T, A/C, or A/G. If the original base call is “A”, then the extra overlapping “A” will create a high “A” peak. The sequencing peaks after the broken template will significantly drop off (DNA run-off). In a partially digested template, some DNA molecules are cut, while others remain intact and the base calls sometimes continue after the sudden peak drop off. Three examples of cut sites from partial digestion are shown near AC N5 GT sites (cleavage taking place either upstream or downstream or on both sides N9-12). Panels 1 and 3 show cleavage upstream at N12 and panel 2 shows cleavage downstream at N9 as indicated by the high “A” peak or a sudden drop in peak height.

#### TRD1 deletion: TRD2-CR2 (no protein expression) and CR1-TRD2-CR2 (nicking)

We attempted cloning and expression of TRD2-CR2 in *E. coli*. The expression plasmid can be constructed, but no protein was detected after IPTG induction and chitin column purification. The reason for TRD2-CR2 poor expression is unknown. However, CR1-TRD2-CR2 protein can be expressed and purified (**suppl. Fig. S2**). This variant generated nicked pBR322 when it was reconstituted with the RM subunits (data not shown). The nicking sites generated by CR1-TRD2-CR2/RM remain to be determined. The 8-bp site (GGAG N5 CTCC) is not present in pBR322, pBC4 (pUC19 with an adenovirus DNA insert, see NEBcutter V3 at NEB.com) or phage λ DNA. Thus, new plasmid DNA substrate needs to be constructed to detect sequence-specific nicking or cleavage.

#### Circular permutation of BsaXI TRDs: TRD2-CR2-TRD1-CR1

It has been suggested that TRDs in a Type I RM system can be shuffled with circular permutation [8]. Due to the modular organization of TRD1 and TRD2, we constructed and purified the rearranged BsaXI TRDs in circular permutated form TRD2-CR2-TRD1-CR1. The S variant is active in cleavage of M13 dsDNA in complex with the RM subunits. **Figure 5** shows the run-off sequencing of two BsaXI sites 5’ GGAG N5 GT 3’ in M13 dsDNA, which confirmed the recognition sequence and slightly shifted cleavage distance at N12-13 and N10-13 over the longer staggered cuts. The shorter staggered cuts were not affected at N7 and N9. It is not clear why TRD2-CR2 variant could not be expressed in *E. coli*, but S variants CR1-TRD2-CR2 and TRD2-CR2-TRD1-CR1 could be expressed and purified.

**Figure 5.**
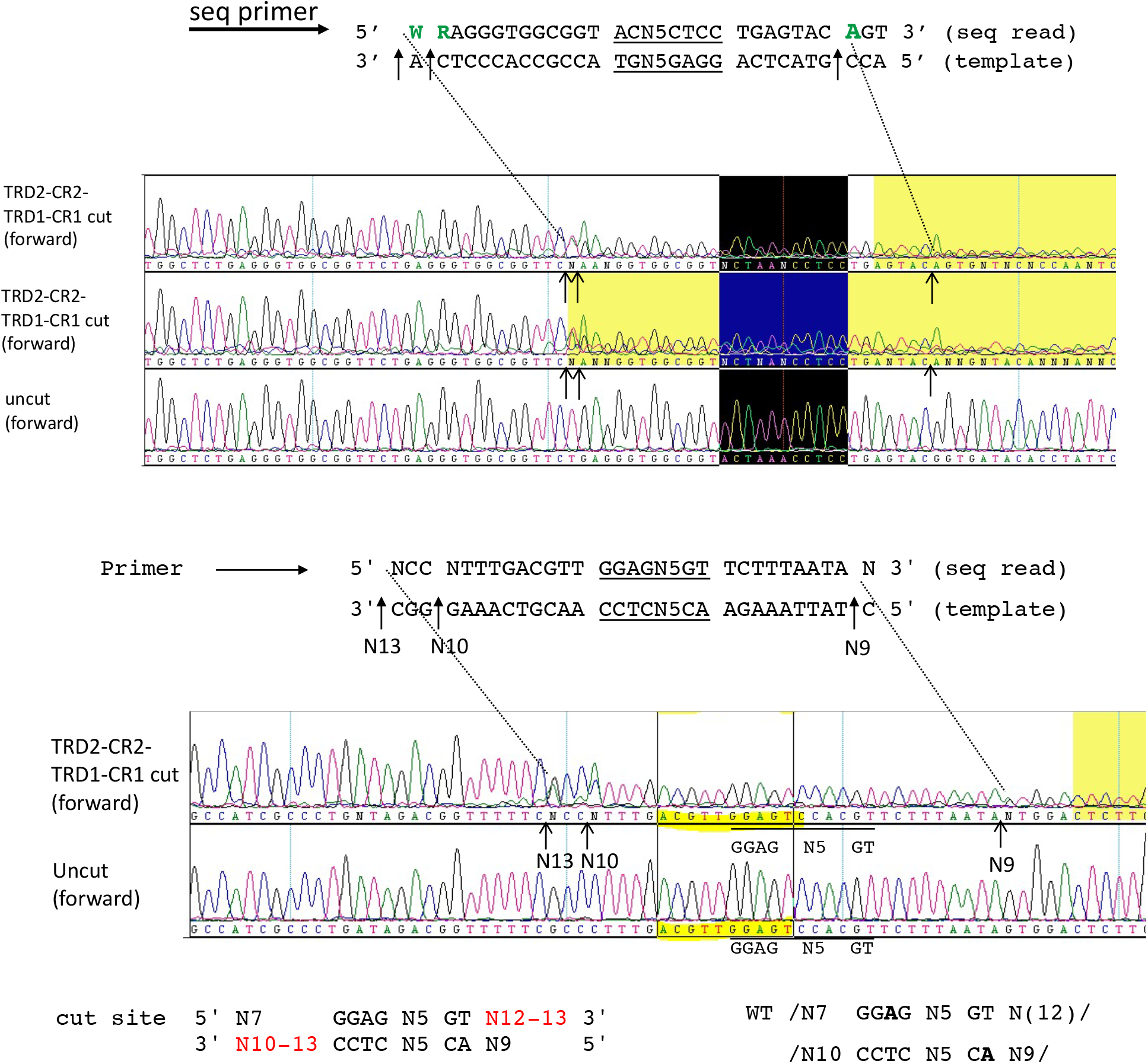
DNA run-off sequencing of digested pBR322 by circular permutated TRD2-CR2-TRD1-CR1/RM. Examples of two cut sites are shown near 5’ GGAG N5 GT 3’. The recognition sequence is identical to the WT enzyme. But the cleavage distance is slightly variable (N12-13 and N10-13). Sequence shaded in yellow: poor sequence read as indicated by the sequence editing software (DNAStar/lasergene). Sequence in the black box indicates the recognition sequence 5’ GGAG N5 GT 3’

#### Replacing BsaXI TRD2 with a TRD homolog with 59% aa sequence identity: TRD1-CR1-TRD2*-CR2

A BsaXI TRD2 homolog CspC0110I TRD (partial sequence) was found in GenBank by a BlastP search. The two TRDs share 59% aa sequence identity (see Materials and Method for aa sequence).

It was annotated as a Type I HsdS partial sequence in *Cyanothece* sp. CCY0110 which is similar to EcoEI HsdS. We constructed and purified the chimeric protein consisting of TRD1-CR1- [CspC0110T TRD]-CR2. Restriction activity was reconstituted when the chimeric S subunit is combined with the BsaXI RM subunits and the cleavage pattern is identical to BsaXI (data not shown). Another S variant (Chimeric S subunit: TRD1-CR1-[Cst TRD]-CR2 (28% aa sequence identity between Cst TRD and BsaXI TRD2) could not be expressed in *E. coli*. We concluded from this experiment that highly homologous TRD (59% sequence identity) can be used to replace BsaXI TRD2 and generate active enzyme with BsaXI RM subunit. TRDs with low sequence identity to BsaXI TRD2 may be problematic possibly due to incompatible TRD with BsaXI CR1 and CR2. The results of all BsaXI TRD rearrangement and mutagenesis are summarized in Table 1.

**Table 1.** Summary of BsaXI TRD rearrangement and mutagenesis. Restriction activity was reconstituted by mixing WT RM subunits with the S subunit variants.

| Specificity (S) subunit | Activity with WT RM | Expression level | Note |
| --- | --- | --- | --- |
| WT: TRD1-CR1-TRD2-CR2 | +++ | + | AC N5 CTCC |
| TRD1-CR1 | ++ | + | AC N5 GT |
| TRD1-CR2 | + (partial) | + | AC N5 GT |
| TRD1-CR1*-8aaΔ | + (partial) | + | AC N5 GT |
| TRD1-CR1*-15aaΔ | + (partial) | + | AC N5 GT |
| TRD1-CR1*-21aaΔ | + (partial) | + | AC N5 GT |
| TRD1-CR1*-32aaΔ | + (partial) | + | AC N5 GT |
| TRD1-CR1*-57aaΔ | NT (not tested) | Poor expression |  |
| TRD1-CR1-TRD1-CR2 (2xTRD1) | + (partial) | + | AC N5 GT |
| TRD2-CR2 | NT | poor expression | ? |
| CR1-TRD2-CR2 | + | + | Nicking activity (Specificity not determined) |
| TRD2-CR2-TRD1-CR1 (circular permutation) | ++ | + | GGAG N5 GT (Imprecise cuts) |
| TRD1-CR1-[CspC0110I TRD]-CR2 (TRD2 replaced by a homolog) | ++ | + | AC N5 CTCC |
| TRD1-CR1-[Cst TRD]-CR2 | NT | poor expression |  |

CR1\*, aa deletion variants in the CR region. CspC0110I TRD, a TRD2 homolog found in the genome of *Cyanothece* sp. CCY0110. Cst TRD, a distant TRD2 homolog found in the genome of *Corynebacterium striatum*. Poor expression means that no target protein of the predicted molecular mass was detected in SDS-PAGE analysis.

#### BsaXI methylase activity

BsaXI is predicted to carry both endonuclease and methylase activity. To detect BsaXI methylase activity of the RM/S complex, phage λ DNA was first incubated with BsaXI in the presence of methyl donor SAM without Mg^2+^ cations, and in the presence of 1 mM EDTA (restriction activity is inhibited by the absence of divalent cations). After two hours methylation, MgCl_2_ (10 mM) was added to the reaction, and 2 U of BsaXI was added to chase the cleavage reaction. After DNA methylation reaction, the DNA was partially resistant to BsaXI digestion (data not shown). The substrate DNA was only partially modified, in agreement with the in vivo expression result that the co-expression of RM and S genes in *E. coli* is toxic and unstable on the same plasmid due to insufficient methylation. It is not clear whether there is another MTase that modified the same sequence from the native host.

#### BsaXI TRD1 and TRD2 homologs: standalone TRDs from sequenced microbial genomes

Only two homologs of BsaXI standalone TRD1 are found in GenBank (WP_052564242=177 aa and KAA0244900=175 aa). Phyre2 search predicted that both TRD1 homologs lack the CR region. Therefore, either sequencing errors resulted in premature termination (missing CR region) or a separate peptide for the CR region or the TRD homologs may function in the absence of CR sequence. In the genome of *Candidatus Brocadia* sinica, two ORFs encoding TRD2 (240 aa) and RM (928 aa) are also present, suggesting they might encode an R-M system (CbrSI). It was noted that Cbr TRD1 and RM putative proteins share high sequence identity to the counterparts in BsaXI (39.3% and 55.9% aa sequence identity). But Cbr TRD2 only shared 23.3% aa sequence identity to BsaXI TRD2. Most BsaXI TRD1 homologs are present as fusion to other TRDs in the S subunits of putative Type I and IIB restriction systems.

BlastP search also identified more than 36 standalone BsaXI TRD2 homologs (196 to 248 aa long). Some homologs are shown in **suppl. Fig. S5**. Most of the BsaXI TRD2 homologs are fused to BsaXI TRD1 homologs or fused to other TRDs with low sequence identity, suggesting BsaXI TRD2 homologs may have partnered with other TRDs to create new specificities.

## Discussion

In this work, we cloned and expressed *bsaXIRM* and *bsaXIS* genes in *E. coli*. BsaXI enzyme was purified by multi-step chromatography from cell extracts containing RM and S subunits and mixed together before purification. We also purified RM and S subunits separately and reconstituted BsaXI activity in vitro by mixing the purified subunits. The reconstituted enzyme activity is lower than the enzyme purified as a natural complex. Similar to BcgI endonuclease complex, BsaXI enzyme also consisted of two RM subunits and one S subunit in the form of [RM]2 S (See **suppl. Fig. S1C and D** for schematic diagrams). The S subunit is analogous to Type I HsdS in domain organization of TRD1-CR1-TRD2-CR2. By rearrangement of TRDs and CRs, we examined the activity of TRD1-CR1, TRD1-CR2, 2xTRD1 (TRD1-CR1-TRD1-CR1) and C-terminal deletion variants in TRD1-CR1 and created a new cleavage specificity 5’ AC N5 GT 3’. As expected, circular permutation of TRDs in TRD2-CR2-TRD1-CR1 created an active enzyme with the same specificity as the WT when the S variant is complexed with RM subunits. The cleavage was not very precise only in the longer nick on the staggered cuts (5’ CTCC N10-13/N7, 5’ GT N12-13/N9), and the shorter nicks at N7 and N9 were not affected. The reason for this different effect is unknown. BsaXI TRD2 domain can be substituted by a homologous TRD protein (59% aa sequence identity) found in *Cyanothece* sp. CCY0110 bacterial genome and the chimeric S subunit can form active complex with BsaXI RM subunits. But a distant TRD homolog with 28% sequence identity failed to express the chimeric S subunit, which may be due to the incompatible protein folding of Cst TRD and BsaXI CR2 (steric conflict). More structure-guided S protein engineering is required to evaluate compatibility between TRDs and CRs. For example, the newly designed TRD-CR protein can be fused to *lacZα* peptide in an *in vivo* protein solubility assay to screen functional TRD-CR protein and its binding activity to cleavage-deficient RM subunit [29].

Our deletion analysis of TRD1-CR1 in the CR1 long α—helix conserved region indicated that the full-length CR1 is not absolutely required for activity, although longer deletion (i.e. 32-aa deletion) had the most impaired activity. The in vitro activity of CR1 32-aa deletion is similar to that of TRD1-CR2 (the entire CR1 was replaced by CR2) with residual partial activity. It is suspected that the spacing distance N5 in the 5’ AC N5 CTCC 3’ bipartite sequence is mostly controlled by the horizontal length of the dumbbell shape of two TRDs (O=O, i.e. the distance of two dumbbells plus the handle) and not entirely determined by the length of the long α—helix conserved regions CR1 and CR2.

There are currently 31 Type IIB restriction systems (REBASE) which cleave DNA sites with bipartite sequences. (http://hae.neb.com/cgi-bin/doublist). Some of them might be amenable to the TRD deletion strategy described here to create new specificities. It is also important to examine the functionality of BsaXI TRD2 specificity. In this work, we demonstrated that CR1-TRD2-CR2 can be expressed and purified. If CR1-TRD2-CR2/RM can form active complex and activate cleavage, the TRD variant may encode a rare cutter (GGAG Nx CTCC). An alternative enzyme discovery strategy is to use BsaXI RM subunit to pair with some BsaXI S homologs with low aa sequence identity (20% to 34% sequence identity) provided that the BsaXI RM subunit can form complex with these homologs to cleave DNA. This screening method may discover new REases with unique but related recognition sequences. In general, Type II REases with more than 35% aa sequence identity encode isoschizomers with the same specificity or with one base off recognition. Enzymes with 22% to 34% sequence identity sometimes encode REases with altered target sites ([7; 30]. More biochemical and structure studies of BsaXI and Type IIB S subunit are needed before we can engineer new enzyme specificities at will from the large number of TRDs found in Type I and IIB restriction systems.

## Acknowledgement

We thank Andy Gardner and Geoff Wilson for helpful discussions and NEB DNA core lab for DNA sequencing. We are grateful to Tom Evans, Richard Roberts, and Jim Ellard for support. Sonal Gidwani was supported by NEB M.S. student internship program. The publication cost was paid for by NEB. Part of this work had been presented at a restriction enzyme conference as a poster presentation.

## Conflict of interest statement

Daniel Heiter and Shuang-yong Xu are employees of New England Biolabs, Inc., a company that develops restriction enzymes and other reagents for research and diagnostic applications.

## Supplementary Material

### Supplement Figures S1-S4

**Suppl. Fig. S1.**
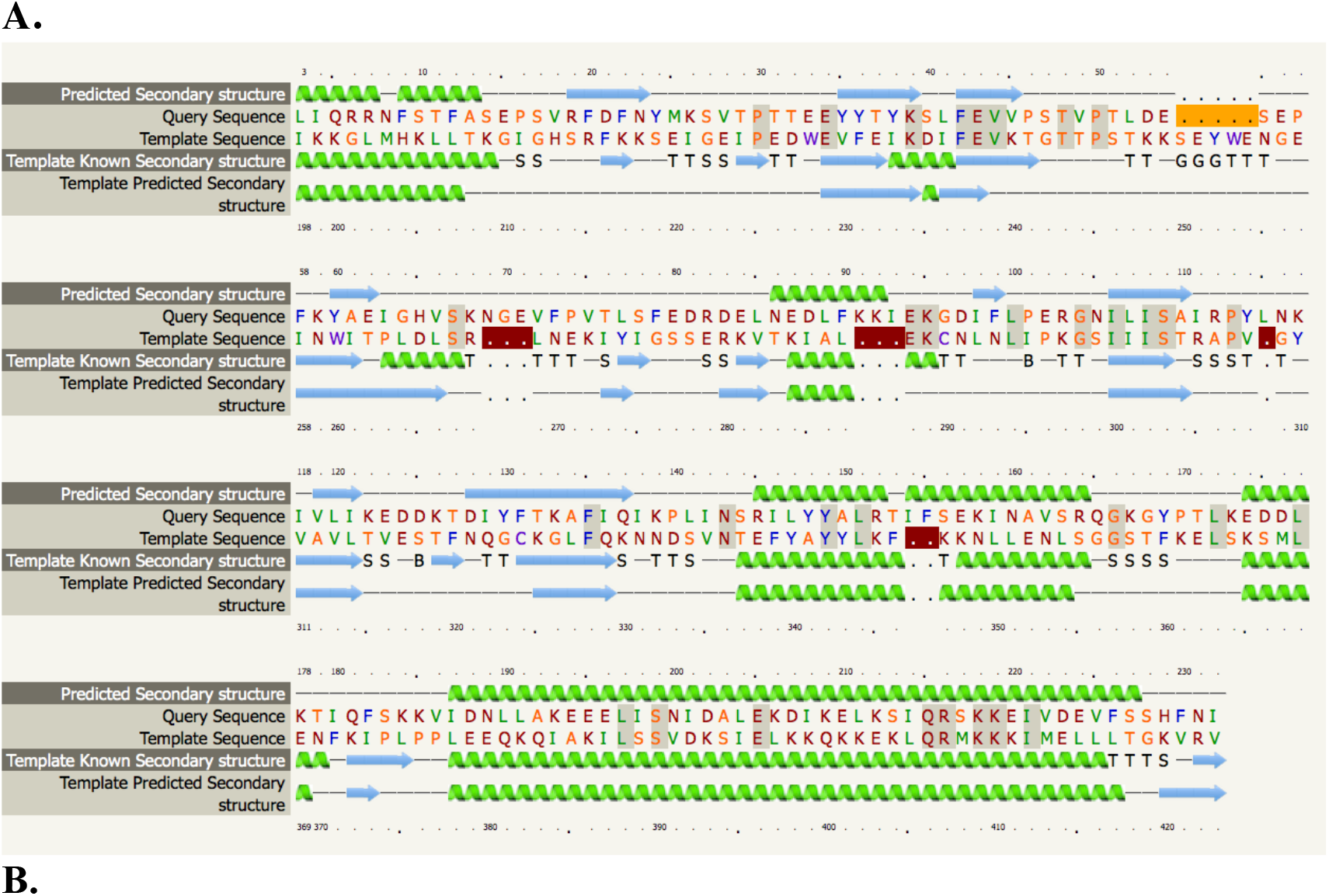
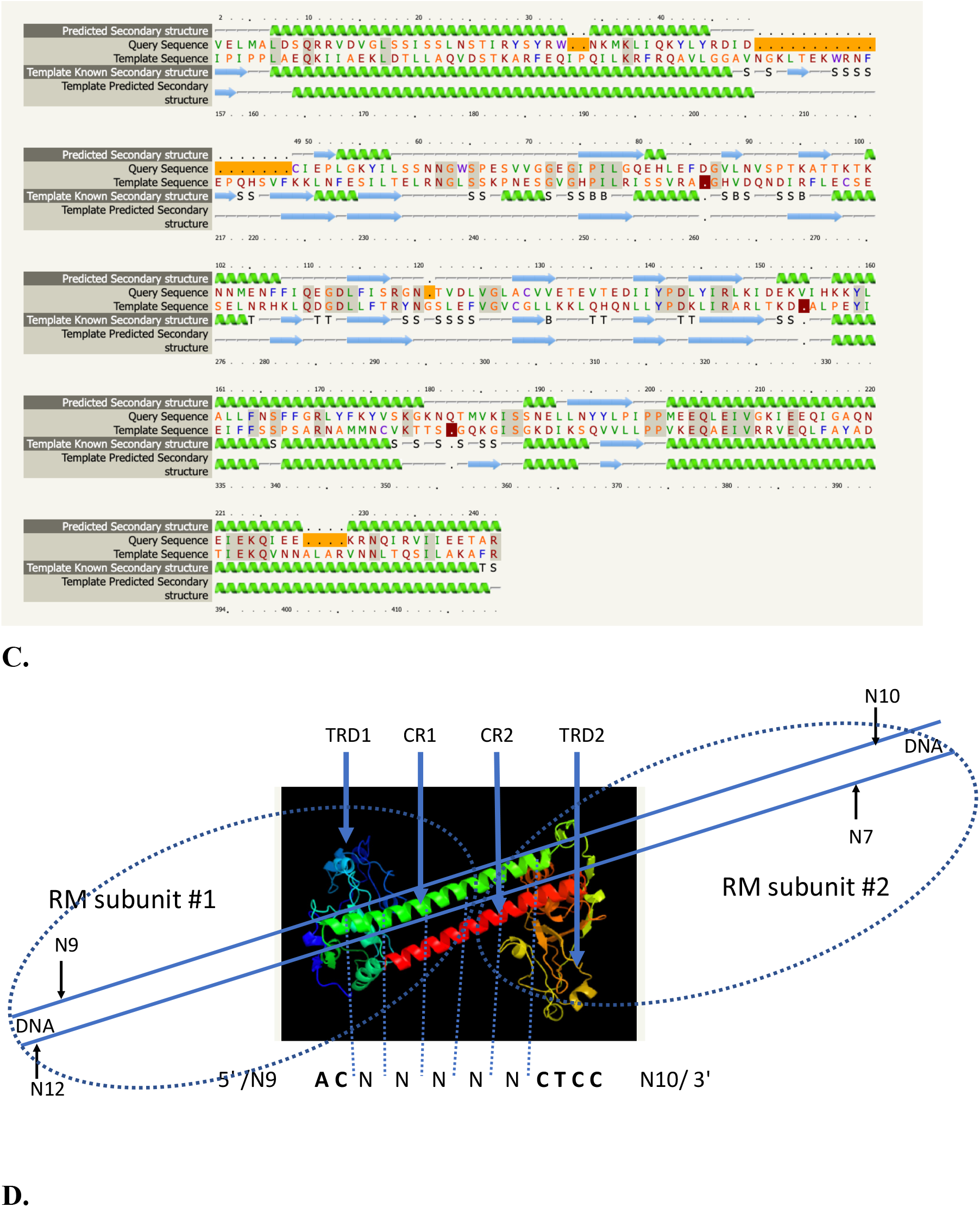
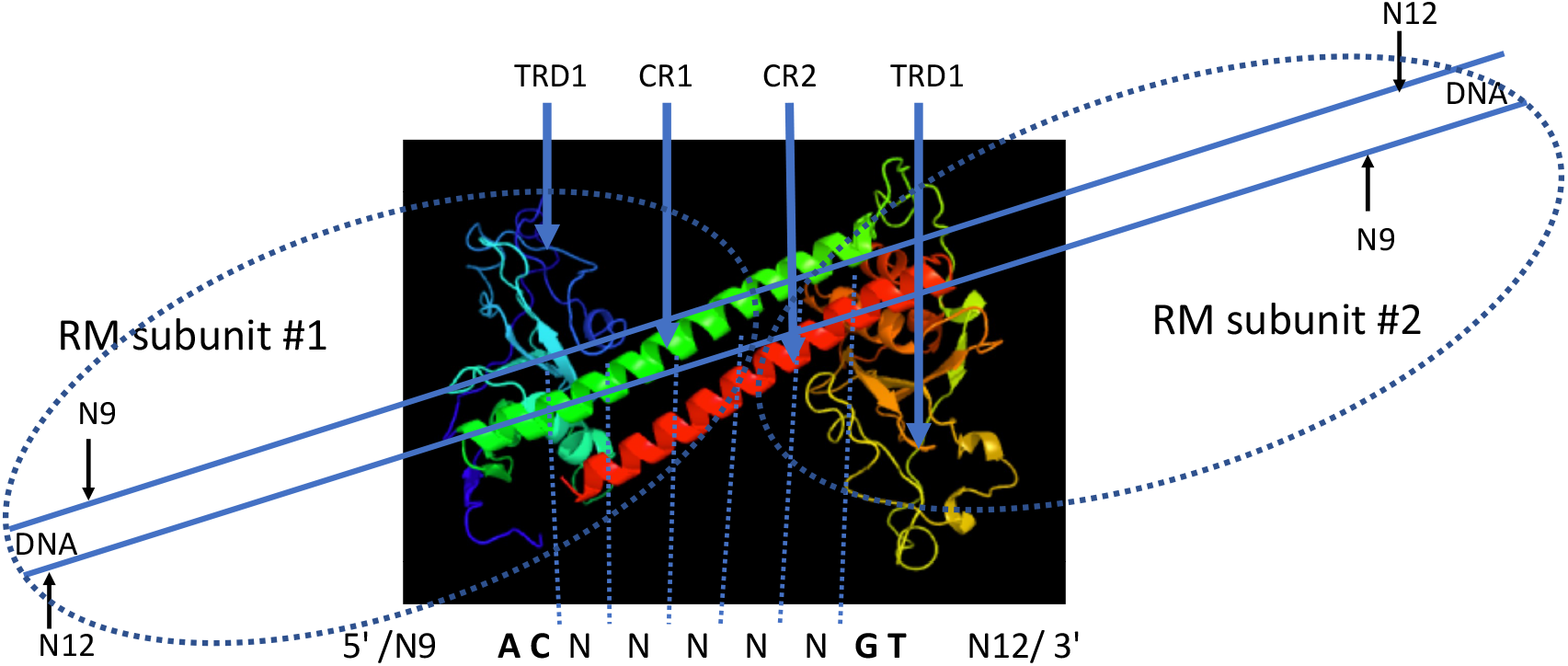
BsaXI TRD1-CR1 (top) and TRD2-CR2 (bottom) predicted secondary structure by the Phyre2. Green coil, α-helix; blue arrows, β-sheets. There are two predicted long α-helixes in TRD2-CR2, located at the N-terminus and C-terminus, respectively. **A and B**, Amino acid sequence alignment of TRD1-CR1, TRD2-CR2 with Type I S subunit TRDs. **C**. BsaXI S structure model predicted by Phyre 2 and schematic diagram of BsaXI S and BsaXI RM subunits. **D**. Predicted 2xTRD1 (TRD1-CR1-TRD2-CR2) model by Phyre2 with schematic diagram of two RM subunits.

**Suppl. Fig. S2.**
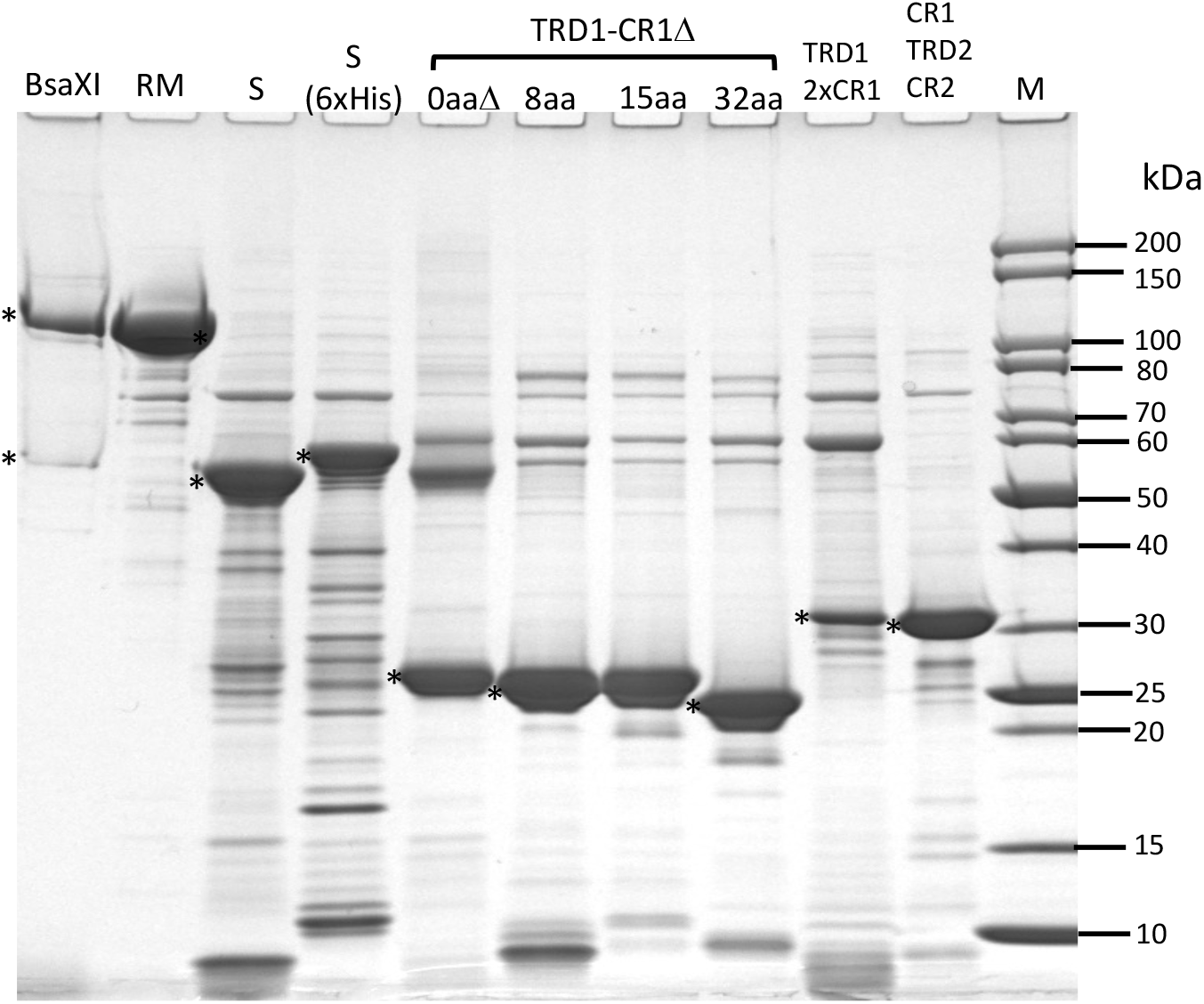
SDS-PAGE analysis of partially purified BsaXI REase, RM, S, S (6xHis) subunits, and S subunit variants. M. protein molecular mass standard in kDa (NEB). Purified target protein is marked by an “*”. BsaXI RM subunit was purified from chitin, DEAE, and Heparin columns; S (6xHis) protein was purified from a Ni agarose column; WT S and S subunit variants were purified from chitin columns by DTT cleavage of target-intein-CBD fusions.

**Suppl. Fig. S3.**
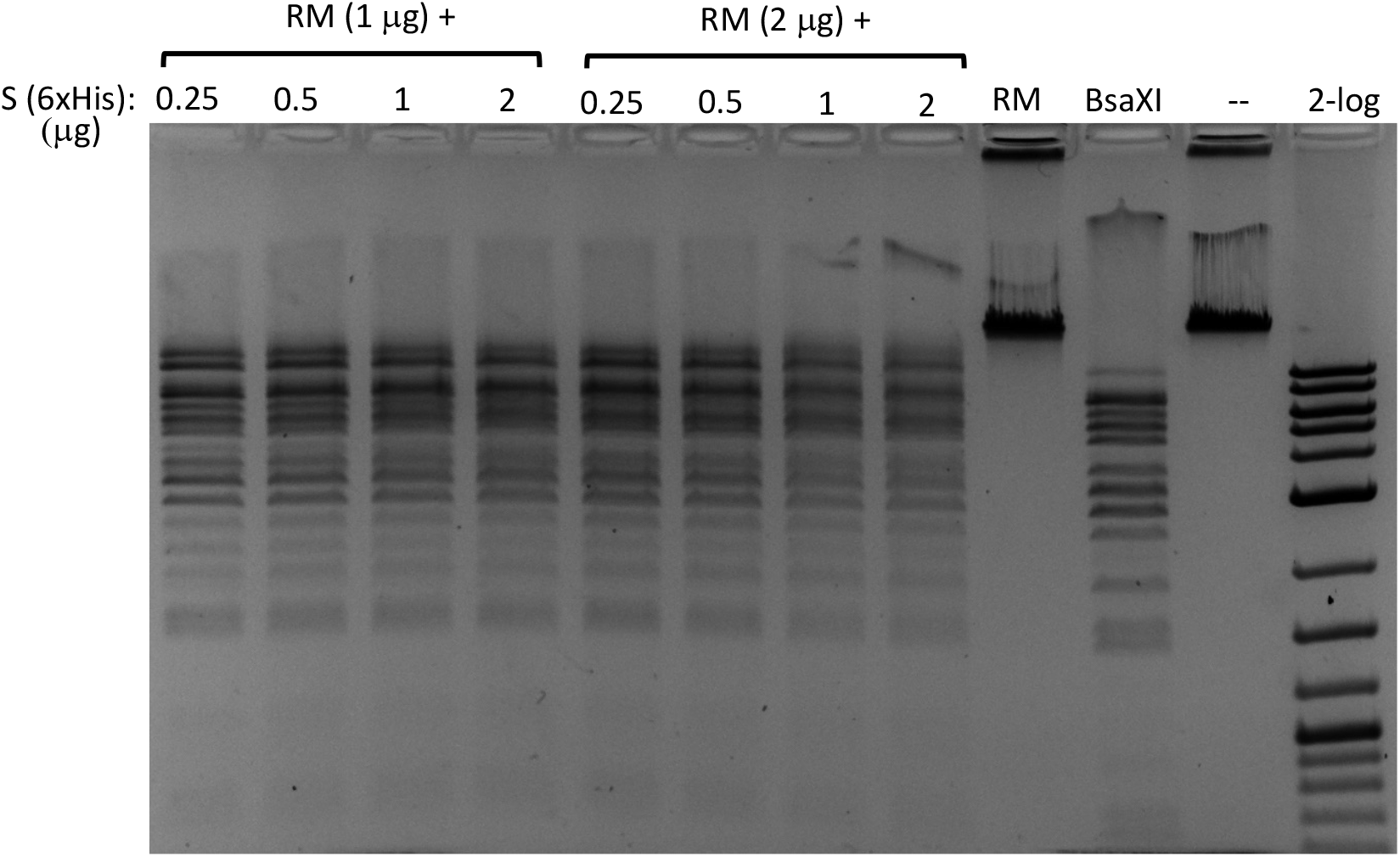
BsaXI restriction activity reconstituted by mixing purified RM (chitin/DEAE/Heparin columns) and S (6xHis) (Ni agarose column) subunits in vitro in restriction digestion of λ DNA. Fixed amount of RM subunit (1 μg, at ∼93.5 nM, lanes 1-4; 2 μg, at ∼187.0 nM, lanes 5-8) was mixed with varying amount of S (6xHis) (0.25, 0.5, 1, and 2 μg, at 90.9, 181.9, 363.6, and 727.2 nM). RM to S ratio approximately at 1/1, ½, ¼, and 1/8 in lanes 1-4; at 2/1, 1/1, 1/2 to 1/4 in lanes 5-8. Lanes 9-11, BsaXI RM subunit only; BsaXI positive control (4 U), uncut DNA. 2-log, DNA size ladder (0.1 to 10 kb, NEB).

**Suppl. Fig. S4.**
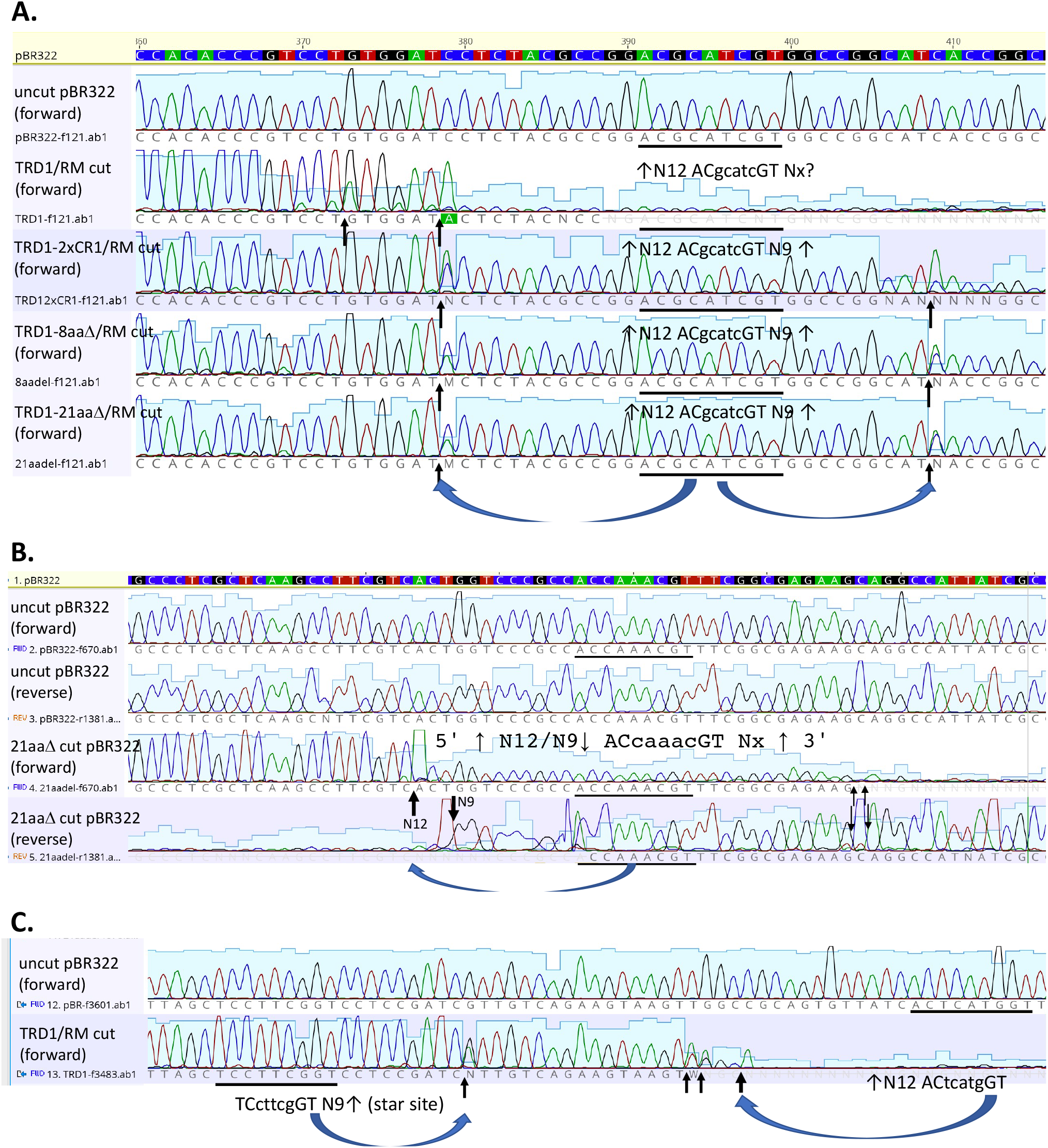
Cleavage site mapping of TRD variants in complex with RM subunits (TRD1-CR1, TRD1-2xCR1, TRD1-CR1-8aaΔ, TRD1-CR1-21aaΔ) by DNA run-off sequencing. **A**. Cleavage upstream (N12) and downstream (N9) of ACgcatcGT site in pBR322 by TRD1, TRD1-2xCR1, and two TRD1-CR1 deletion variants (sequencing panels 2 to 5) compared to undigested DNA (sequencing panel 1). **B**. Cleavage upstream of ACcaaacGT site by TRD1-CR1-21aaΔ. Cleavage took place mostly upstream and a low level of cleavage downstream as the result of asymmetric partial digestion. **C**. Possible cuts downstream of a star site TCcttcg GTN9↓ and cleavage upstream of a cognate site ↓N12ACtcatgGT.

**Suppl. Fig. S5.**
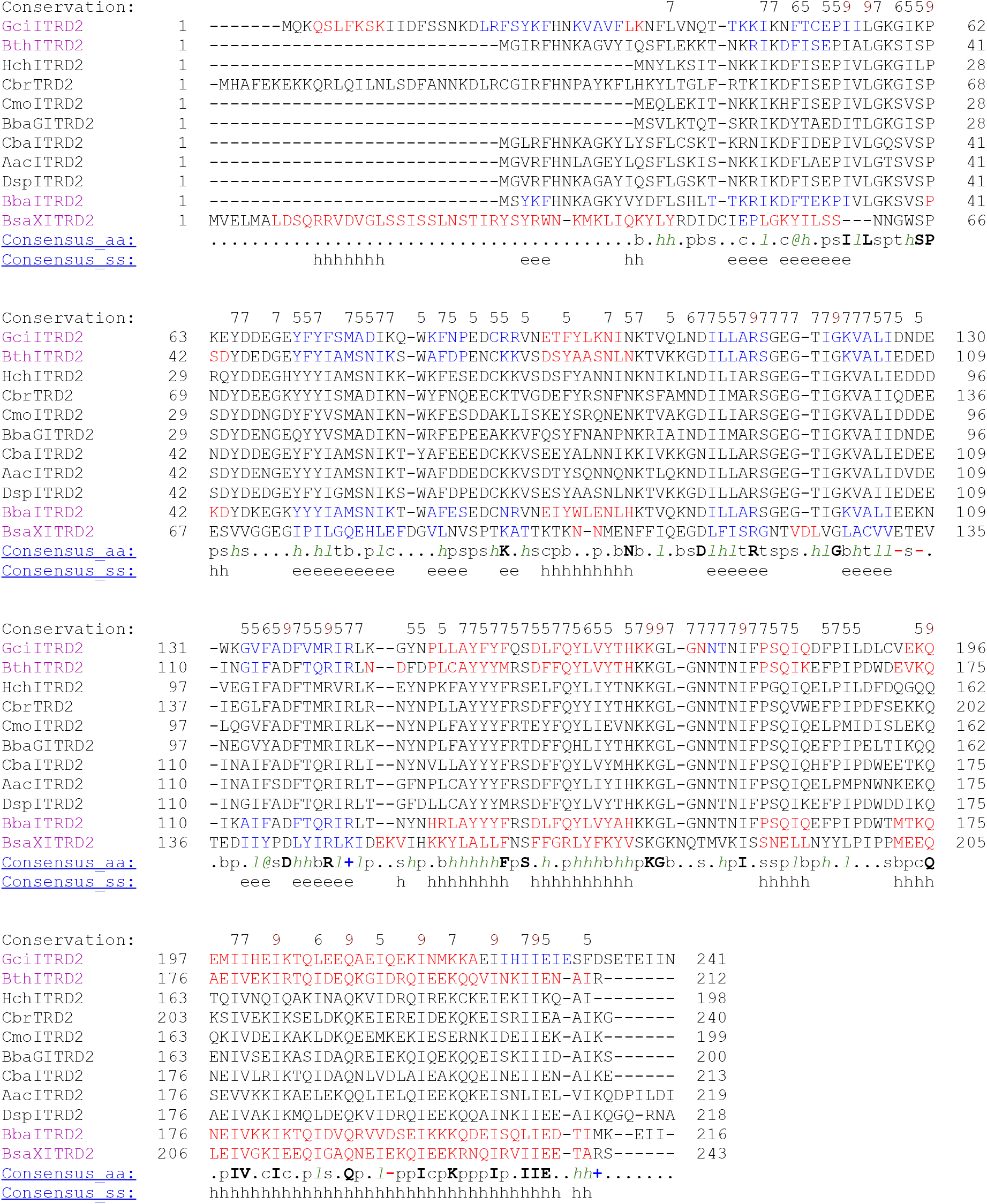
PROMALS3D multiple sequence alignment of 10 standalone (orphan) TRDs (BsaXI TRD2 homologs) from sequenced microbial genomes. The numbers (5-9) on top of aa sequences indicate the degree of conservation at the particular positions. Predicted α-helix = h; predicted Δ-sheet = e. The highly conserved aa residues are shown in bold letter. There are nine conserved Ile (I) and Leu (L) residues in the C-terminal CR2 region that are presumably important for interactions with CR1 and cognate RM subunits to form helix bundles.

